# FlAsH-ID: A short peptide tag for live cell photoproximity labeling

**DOI:** 10.1101/2025.10.24.684417

**Authors:** Wuyue Zhou, Shashank Nagaraja, Jihye Kim, Thomas Venables, Matthew E. Pipkin, Ciaran P. Seath

## Abstract

Methods to determine how biomolecules interact in a cellular context has led to significant advances in our understanding of biology. Of these, proximity labeling has emerged as a valuable tool for the interrogation protein-protein and protein-RNA interactions. Almost all proximity labeling methods are deployed via large fusion proteins, which while powerful in many contexts, can disrupt native biology or obscure important epitopes on the protein under study. This challenge is especially prominent for small transcriptionally active proteins where the protein cistrome is vital for correct localization and function. Here, we have developed a photoproximity labeling method that is localized to a short 6 amino acid peptide sequence that employs the combination of the Tsien tetracysteine tag with FlAsH dye for localized singlet oxygen generation that we term FlAsH-ID. We demonstrate this method is non toxic and can be employed to label several cellular compartments, and generate high quality interactomics data around H3.1. Furthermore, we show that FlAsH-ID can be used to probe dynamic chnages in protein movement in response to immune stimuli, identifying several RNA-binding protiens that are trafficked to the nucleus following poly(I:C) treatment. Finally we demonstrate FlAsH-ID can be used for proximity proteomics in primary CD8^+^ T cells by focusing on the transcription factor RUNX3. RUNX3-tetracysteine, but not RUNX3-TurboID, recaptiulated the function of wildtype RUNX3 and rescued the phenotype of *Runx3-*deficeint CD8^+^ T cells in response to viral infection *in vivo*. In primary CD8^+^ T cells, FlAsH-ID identified RUNX3 in proximity to the nucleosome remodeler ARID1A and we show that both factors act concertedly during initial naive CD8^+^ T cell activation to drive chromatin accessibility in *cis*-regulatory regions that induce differentiation of effector and long-lived memory CD8^+^ T cells. Our results suggest this technique will expand the field of proximity labeling to small, functionally sensitive proteins in primary cells without perturbing their function.

## Introduction

Proximity labeling has emerged as a valuable tool for interrogating interactions between proteins and RNA, being employed widely across the biological sciences^1,2^. These tools have found particular utility in the study of insoluble proteins (e.g., chromatin, cell membrane) where co-immunoprecipitation fails to capture native interactions due to the harsh conditions required to solubilize DNA/protein and lipid/protein interactions^3,4^.

The most commonly used methods remain the enzyme fusions APEX and BioID, and their derivatives, which can be readily inserted into any protein of interest^5^. While powerful and broadly applicable, the fusions required for these techniques are relatively large (∼30kDa), which can perturb native biology, particularly when attached to small proteins, peptides, and proteins where the termini engage in specific biological processes^6–8^. For practitioners studying such proteins, there are no options available for the generation of interactomics data in a native environment (Fig. 1a).

**Fig 1.**
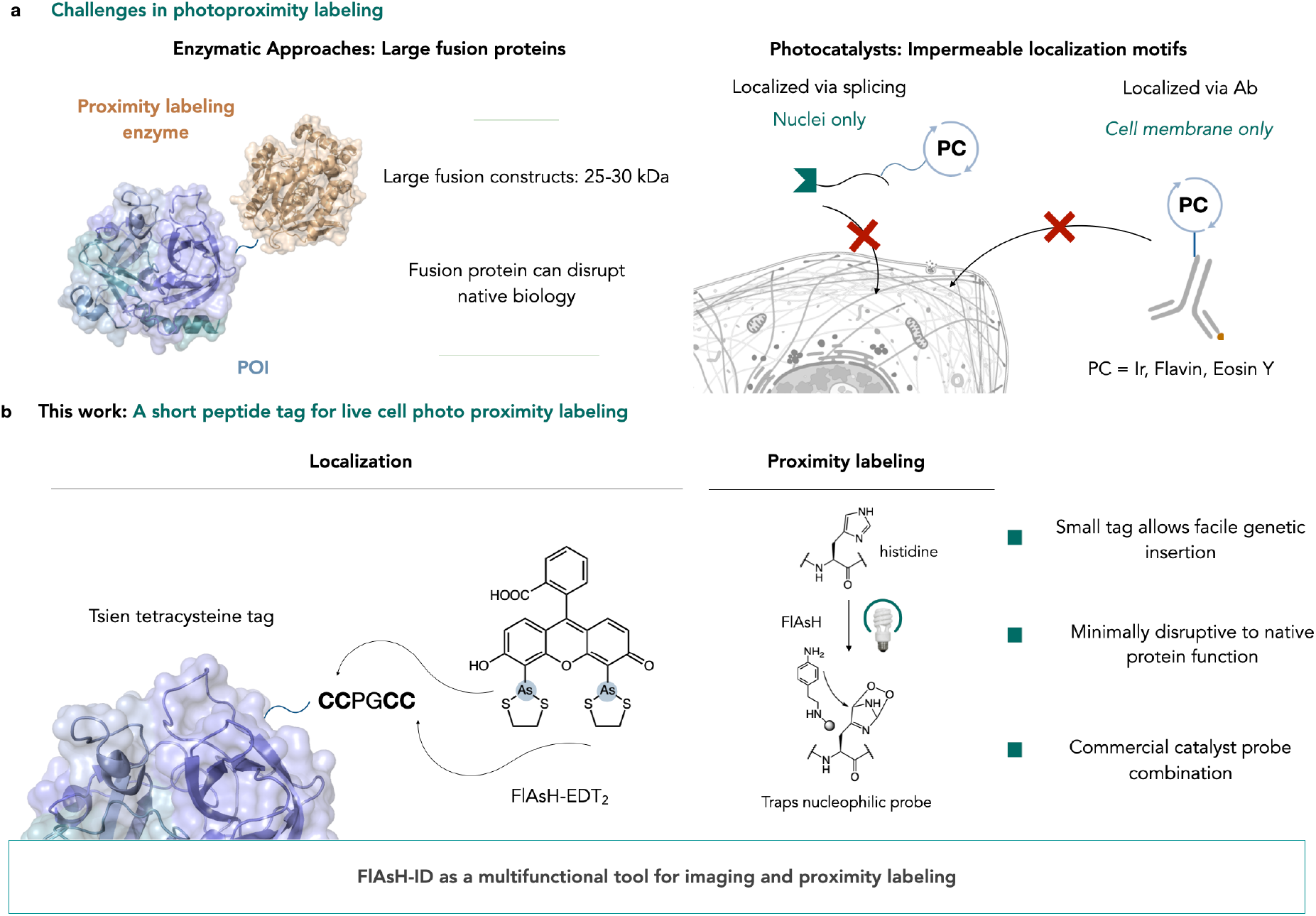
Development of FlAsH-ID: A photoproximity labelling tool that requires only a short peptide insertion. **a**, Current limitations faced by common proximity labeling tools. Genetically encodable enzymatic approaches are easy to administer but the insertion of bulky fusion proteins may perturb protein function. Photocatalyst-based approaches avoid gene manipulation but requires antibodies or fusion proteins to localize to POI. **b**, FlAsH-ID scheme. The dye FlAsH-EDT_2_ has been repurposed to act as a photocatalyst. Following administration, it localizes to the 6-amino-acid-long tetracysteine tag conjugated to the protein of interest, and labels surrounding neighborhood under light stimulation by generating singlet oxygen species which oxidize histidine residues to peroxides that are trapped by supplied nucleophilic probes.

Recent advances in the field have moved towards the use of visible light to initiate labeling, providing fine temporal control and modularity in probe and catalyst construction that can be tailored to each use case^9^. Most of these methods have been deployed extracellularly through small molecule photosensitizers appended to antibodies for cell surface labeling while fewer methods exist for intracellular labeling as incorporation of photocatalysts to specific cellular locations represents a significant challenge^10-12^ (Fig. 1a). Despite this, when compared to enzymes, photocatalysts are able to initiate proximity labeling with a relatively small footprint (<1kDa).

Inspired by these approaches, we sought to develop a method that incorporated the advantages of both approaches. i.e., a photocatalytic method that is minimally disruptive to the native environment and can be deployed to any cellular location. Additionally, an ideal method would employ a short epitope tag that can be readily integrated into either plasmid DNA or directly into the genome via gene editing.

Such a proposal would require a peptide specific chemical ligation that can be performed in live cells. To the best of our knowledge only one method for this has been reported, the FlAsH-EDT_2_ (FlAsH) method and its derivatives from the Tsien group^13^. Here, a tetracysteine motif (CCPGCC) is incorporated into a protein sequence, which can then bind a derivatized fluorescein molecule in live cells^14^ (Fig. 1b). This method has been used for both imaging and stoichiometric crosslinking across a variety of cellular contexts^15–18^.

## Method development

We questioned whether we could adapt this localization strategy for photoproximity labeling, combining the advantages of FlAsH localization with the temporal control of photoproximity labeling, a method we term FlAsH-ID. We reasoned that FlAsH might be able to initiate proximity labeling through either single electron transfer or generation of singlet oxygen. We began our studies by irradiating HEK293T cell lysate for 10 mins with 450 nm light with a series of biotinylated probe molecules (Biotin-Diazirine, Biotin-Aniline, Biotin-Azide, and Biotin-Phenol) in the presence of or absence 2.5 µM FlAsH (Fig. 2a).

**Fig 2.**
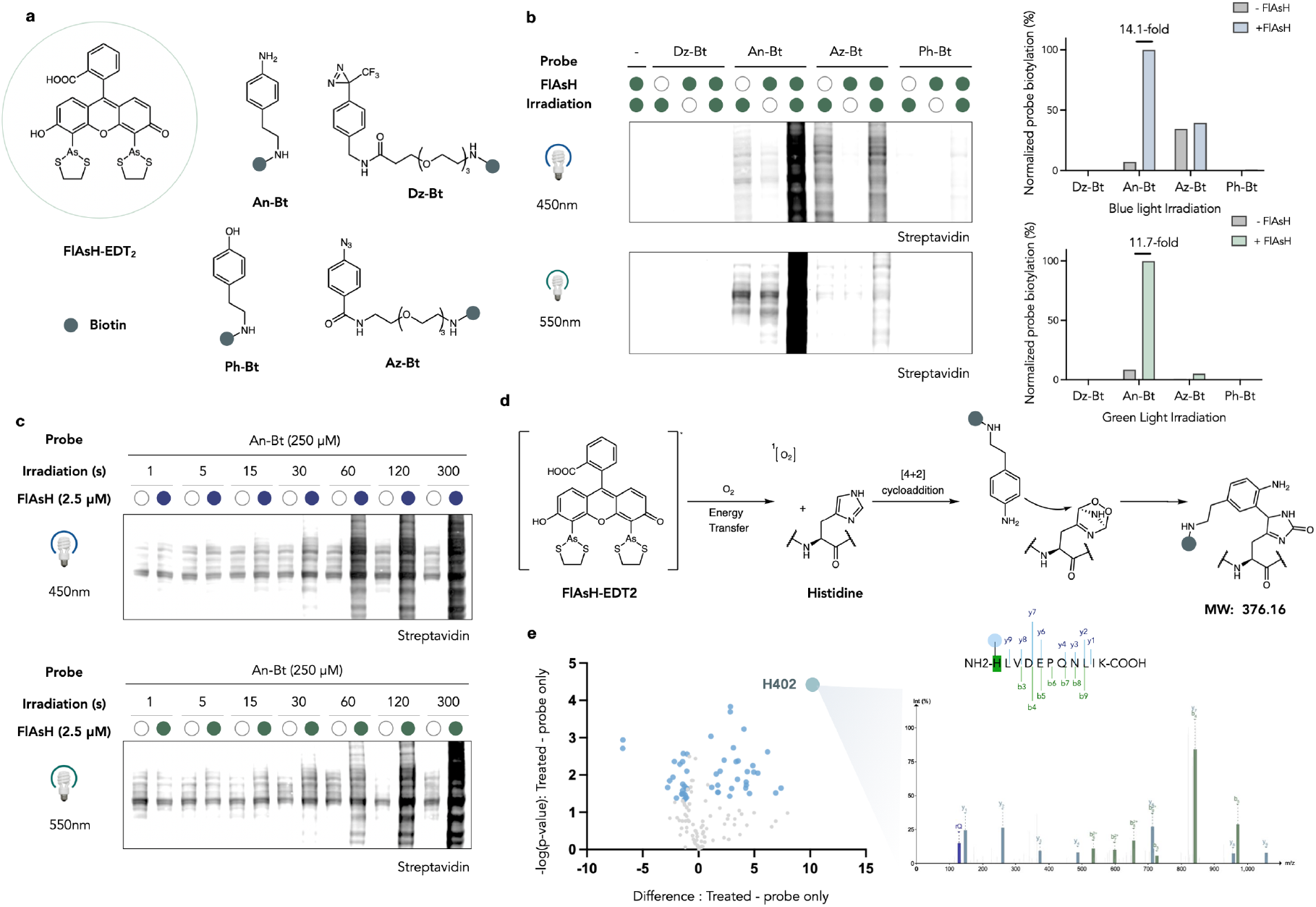
Repurposing FlAsH-EDT_2_ for photocatalysis and development of a proximity-labelling platform. **a**, Structure of catalyst and probes screened in this study. **b**, Screening biotin bearing probes with cell lysate. In the presence of aniline-biotin, cell lysate has been biotinylated only following catalyst and light administration. Comparable labelling is generated by blue and green light, as shown by western blotting. **c**, Time-dependent activation of aniline-biotin probe with FlAsH-EDT_2_. 5 min irradiation with green light gives optimal excitation. **d**, Proposed mechanism of FlAsH-EDT_2_ activation. Following light administration, generated singlet oxygen locally oxidizes histidine residues, which forms endoperoxide intermediates that are trapped by the nucleophilic aniline probe and causes a mass shift of 376.16 Da on the histidine residue. **e**, Volcano plot derived from a two-sided *t*-test showing bovine serum albumin (BSA) peptides enriched in a comparative proteomics study between FlAsH-activated BSA and no catalyst control. Blue dots represent significant enriched peptides. Illustrative MS2 spectra has been presented for peptide HLVDEPQNLIK showing modification of histidine 402 with the desired mass shift.

We observed no labeling with the diazirine probe, which was expected based on the triplet energy of the parent fluorescein dye (Fig. 2b). Conversely, biotin-aniline was extremely reactive under these conditions, showing 14:1 signal to noise versus a control without FlAsH and 59:1 vs a control without light. Under blue light irradiation, biotin-azide showed significant background reactivity, providing little evidence for productive catalysis. Finally, biotin-phenol showed minimal labeling over background. This observation is consistent with a mechanism driven by singlet oxygen generation (SOG) compared to single electron transfer (SET), although SET does appear to be operable with the azide and phenol probes, but dramatically less efficient. Interestingly, performing the same experiment with green light irradiation led to a small decrease in protein biotinylation (12:1 signal to noise ratio) with biotin-aniline, but eliminated most of the background labeling with biotin-azide, providing a 2:1 signal to noise ratio. We next demonstrated that the reaction was dependent on irradiation time, probe concentration, and catalyst concentration (Fig. 2c, supplementary Fig. 1a, b). These data suggested 5 mins with 2.5 µM catalyst and 250 µM probe as a useful set of conditions to move forward with.

To prove our mechanistic hypothesis (Fig. 2d), we used our approach to label recombinant BSA and analyzed tagged peptides by mass spectrometry and found only a single site of labeling at a surface exposes histidine residue (His402) being enriched with a mass shift consistent with the SOG mechanism rather than SET (Fig. 2e, supplementary Fig 1c, d). Furthermore, the reaction was effective to a lesser extent with biotin-hydrazide, a probe known to be compatible with SOG, and the labeling signal was ablated following addition of singlet oxygen quenchers (supplementary Fig. 1e).

We next turned our attention to the development of a protocol for live cell proximity labeling (Fig. 3a). Cognizant of the potential for cellular toxicity derived from the FlAsH catalyst, we treated HEK293T cells with either fluorescein or FlAsH for either 1 or 48 hours to understand the impact of the catalyst on cellular fitness (Fig. 3b, left panel). While fluorescein was tolerated at all concentrations over 48 h, FlAsH led to cell death at micromolar concentrations after 48 h but was non-toxic after a one-hour incubation even at high concentrations. Treatment for 1 h with FlAsH followed by washing steps typically associated with FlAsH fluorescence imaging led to no observable toxicity after 24 h, suggesting that once localized, FlAsH is minimally toxic and could be deployed in a cellular context for extended periods of time without concern (Fig. 3b, right panel). Taken together, we were confident that we could perform proximity labeling experiments without dramatic changes to cellular physiology.

**Fig 3.**
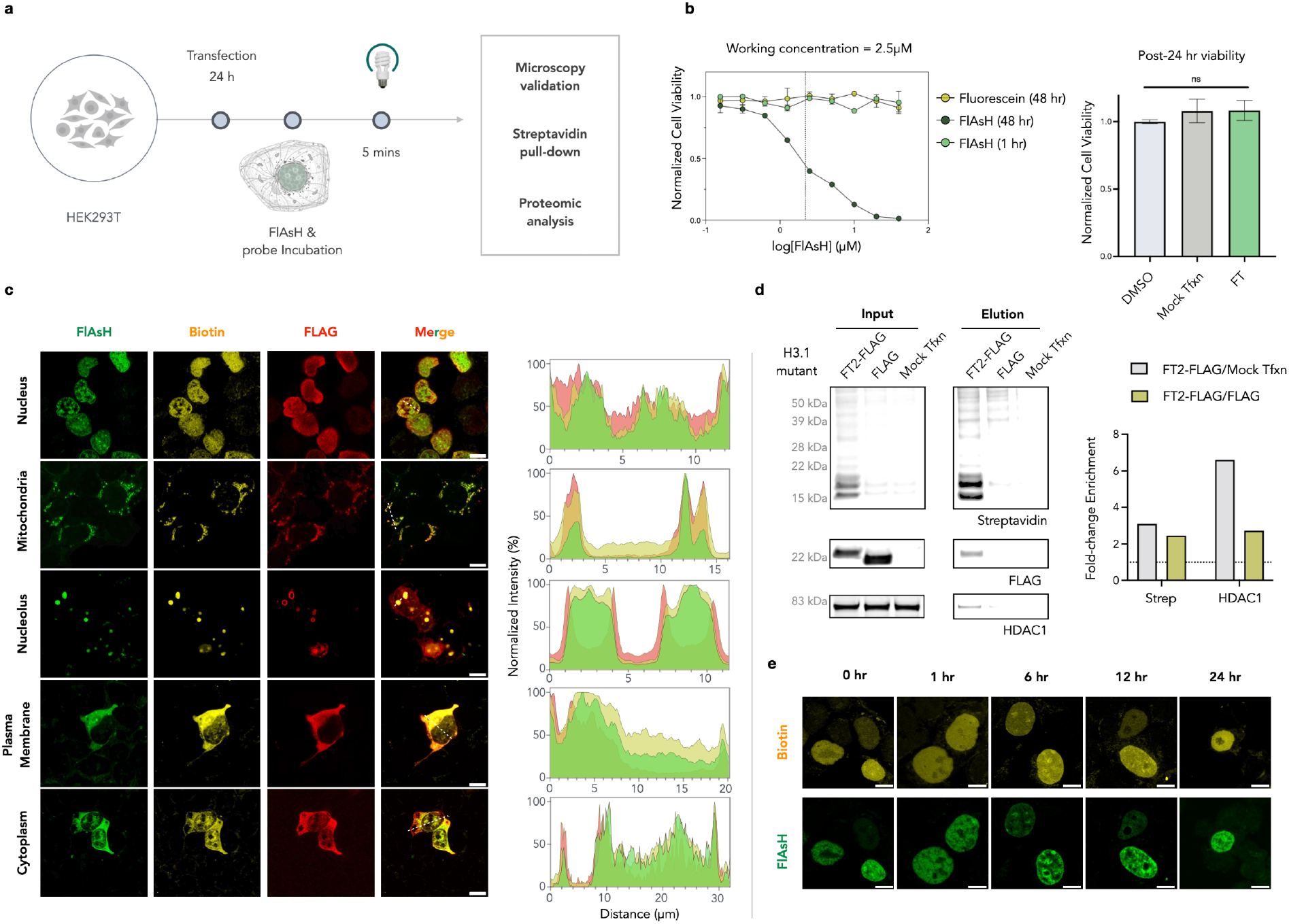
FlAsH-ID displays excellent cellular compatibility and is retained in cells. **a**, Cartoon of FlAsH-ID administration in a cellular system. **b**, Toxicity of FlAsH-ID in cells. Left, prolonged treatment of the catalyst leads to cell death but minimal toxicity is observed with 2.5 µM working concentration over 1 hr (mean with SEM, *n*=3 independent biological replicates, IC_50_=2.21 µM for 48 hr treatment, calculated with nonlinear regression). Right, post-24 hr cell viability is not affected by performing FlAsH-ID (mean with SEM, *n*=4 independent biological replicates, one-way ANOVA, *P*<0.05). **c**, Colocalization of biotinylated signal, target protein, and FlAsH dye in different cellular compartments via immunofluorescence assay. Protein constructs generated for specific compartments are listed in text. Scale bar = 10 µm. **d**, Validation of FlAsH-ID using histone H3.1 with tetracysteine tag addition (H3.1-FT2), in nuclear fraction H3.1-FT2 has been significantly enriched after streptavidin pull-down, as shown by western blotting. **e**, Incorporated FlAsH dye and biotin signal is sustained in the cell for up to 24 hr.

### Proximity labeling in live cells

We began our protocol development by expressing H3.1 with a C-terminus CCPGCC (FT tag) or a longer HRWCCPGCCKTF tag with higher affinity (FT2 tag) flanked by FLAG in HEK293T cells to assess FlAsH localization and labeling specificity^19^. We found that we could effectively localize FlAsH to chromatin by incubation with 2.5 µM for 0.5 h followed irradiation with green light yielded strong nuclear biotin signal by confocal microscopy that was only observed when all components (light, FlAsH, probe, H3.1-FT2) were present (supplementary Fig. 2a). Assessment of biotinylation by western blot showed a requirement for all components and demonstrated a significant enrichment in both the bait and true positive interactor, HDAC1, which were only pulled down when all components were present (Fig. 3d).

We next tested the stability of the incorporated FlAsH labeled cells, by treating and measuring nuclear fluorescence over 24 h. Pleasingly, nuclear fluorescence was maintained over this time course with no observable toxicity, suggesting that this protocol can be used to study cellular stimuli that occur over longer timeframes (Fig. 3e).

To test the generality of our approach we incorporated the tetracysteine tag onto proteins localized to different cellular compartments and assessed localization and labeling specificity by confocal microscopy. Catalysts incorporated in the nucleus (H3.1), nucleolus (3 tandem nucleolar targeting sequences from NIK protein), cytosol (HINT1), mitochondria (PDK1), and cellular membrane (B2) were all well tolerated with this protocol, providing localized biotinylation and fluorescence (Fig. 3c). Aligned with the western blot data, the biotinylation signal is proportional to the amount of light and probe added (supplementary Fig. 2b). Importantly, incorporation at both C- and N-termini of H3.1 were equally well tolerated (supplementary Fig. 2c).

We next established a tandem mass tag (TMT)-based quantitative chemoproteomics workflow to determine the interactome of H3.1 using our approach (Figure 4A). We labeled HEK293T cells transfected with H3.1-FT2 for 5 and 10 minutes and compared to an untransfected control (3 biological replicates per condition) (Fig. 4a). Each condition was lysed and enriched on streptavidin beads prior to on-bead digestion and TMT labeling. Pooled samples were reverse phase fractionated into 10 fractions and analyzed by liquid chromatography-tandem mass spectrometry (LCMS/MS). We identified 7912 proteins from this experiment (supplementary Fig. 3a) with strong correlation between experimental replicates, as shown by PCA analysis (supplementary Fig. 3b). Receiver operating characteristic curves were plotted to determine TMT-ratio cut offs for positive vs negative comparisons, using the mitochondrial proteome as true negatives and BIOGRID interactome as true positives^20,21^. This analysis demonstrated that the accuracy of our approach was comparable to existing PL methods such as APEX, as it gave a smooth ROC curve and the area under curve (AUC) value reached 0.88 (supplementary Fig. 3c). A cut-off value of the log_2_FC ratio was calculated to be 0.142, by maximizing the difference between TPR and FPR to denoise the proteome. 1651 proteins met this cut-off in addition to meeting statistical significance (FDR<0.05) (Fig. 4b). Analysis of detected proteome showed that our method was able to capture 52% of the known nuclear proteome and 90% of all known interactors of H3.1 (BioGRID) (Fig. 4c). Furthermore, of the 1651 enriched hits, 75% were annotated as nuclear by human protein atlas. Of the enriched hits, nucleosomal components comprised 6 of the top 12 hits (log_2_FC > 1.5). Gene ontology analysis of the interactome was enriched in chromatin-based terms such as DNA replication (FDR=9.32E-22) and chromatin organization (FDR=1.03E-71) (Fig. 4d). We observed a similar trend of enrichment for nucleosomal components after performing the labeling reaction with 10 mins of green light irradiation, with many overlapping hits (Supplementary Fig. 3d, e). FlAsH-ID with FT tag on the N-terminal histone tail was also able to enrich the bait and nucleosomal components as top hits, however, with a lower coverage of the nuclear proteome (Supplementary Fig. 3f).

**Fig 4.**
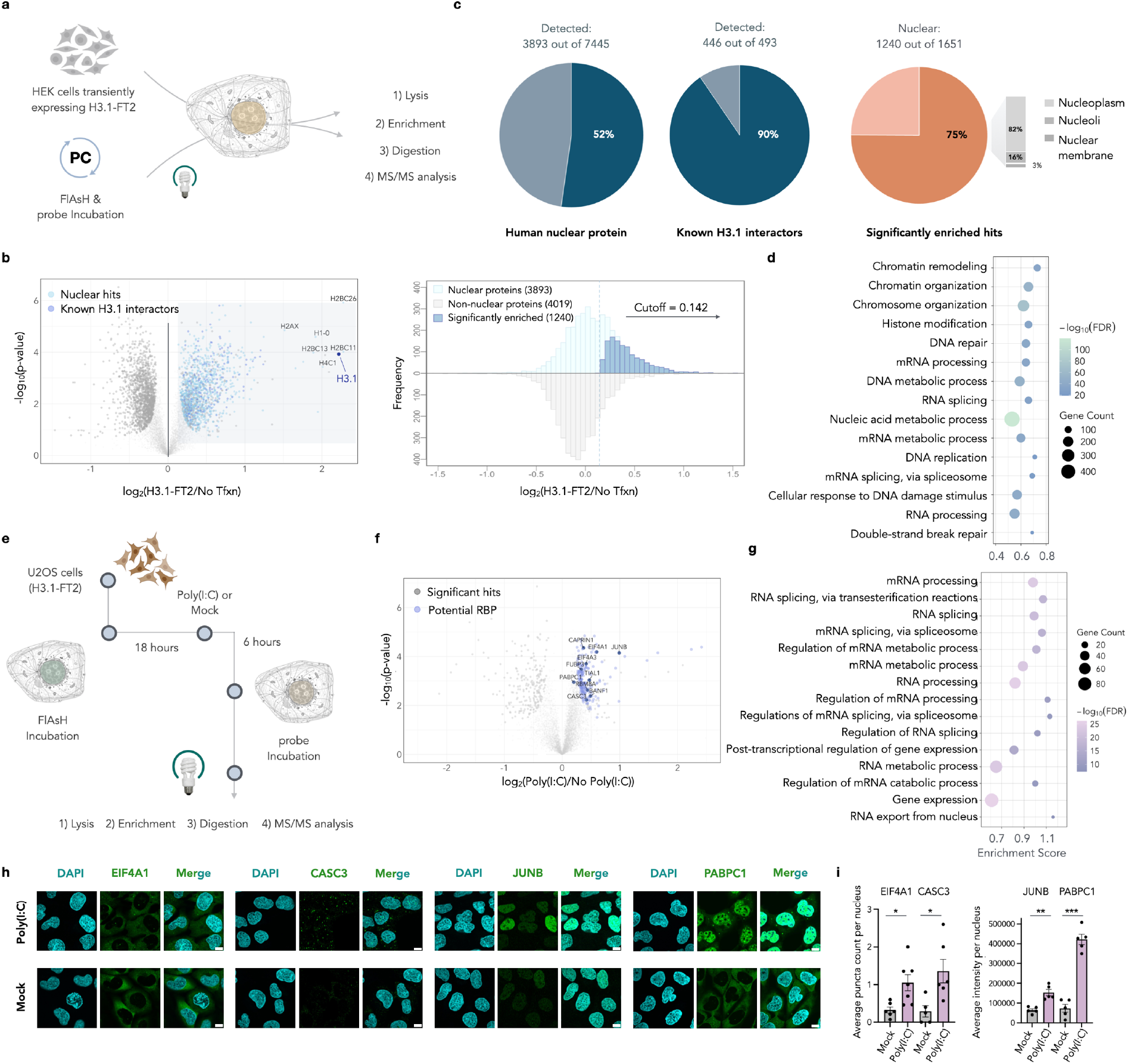
Assessing static and dynamic protein interactomes with FlAsH-ID. **a**, Scheme of a general FlAsH-ID LC/MS workflow. H3.1-FT2 was transiently expressed in HEK293T cells. **b**, Dataset showing protein distribution. **c**, A volcano plot derived from a two-sided *t-*test showing H3.1 interactors from the FlAsH-ID method comparing to untransfected HEK cells. Nuclear proteins, known interactors, and other nucleosome subunits are highlighted. Cutoff was determined by ROC analysis and thresholded with FDR<0.05. **d**, Histogram distribution of log_2_(FC) for nuclear and non-nuclear proteins, derived from the H3.1-FT2 interactome data. Cutoff was determined by ROC analysis. **e**, GO biological processes analysis for H3.1-FT2 versus non-transfected cells. Terms are ranked by a weighted harmonic mean between the enrichment score and -log_10_FDR. **f**, Scheme of tracking dynamic changes in chromatin interactome with FlAsH-ID. U-2 OS cells stably expressing H3.1-FT2 were preincubated with FlAsH 18 hr prior to poly(I:C) which is used to mimic viral infection. Trafficking proteins were labelled following An-Bt and green light administration. **g**, A volcano plot derived from a two-sided *t-*test showing H3.1 interactors from FlAsH-ID analysis comparing poly(I:C) treatment and mock transfection. Potential RBP are highlighted and known and novel RBP targets are annotated. **h**, GO biological processes analysis for proteins with increased abundance in the nucleus. Scale bar = 10 µm. **i**, Validation of hits trafficking into the nucleus following poly(I:C) treatment via immunofluorescence assay (mean with SEM, *n*>=5, two-sided *t-*test were performed for statistical significance. \**P*<0.05, \*\**P*<0.01, \*\*\**P*<0.001).

### FlAsH proximity labeling measures changes in protein localization

Confident our approach can accurately determine nuclear interactomes, we wanted to assess our ability to measure changes to this interactome in response to external stimulus. RNA-binding proteins (RBP) are known to traffic to the nucleus in response to dsRNA lipofection, however an unbiased analysis of protein movement in native environment has never been reported^22,23^. To assess this, we treated U-2 OS cells stably expressing H3.1-FT2 with FlAsH for 0.5 h, and washed with excessive EDT to provide precise incorporation of FlAsH within the nucleus (Fig. 4e, supplementary Fig. 4a). We then treated with 1 µg of poly(I:C) per 1M cells or mock for 6 h to initiate the immune response and movement of RBPs to the nucleus followed by FlAsH-ID labeling and MS/MS analysis (supplementary Fig. 4b). We observed 229 significant proteins that increase abundance in the nucleus in response to poly(I:C), of which 190 are annotated RBPs according to the RBP2GO database (Fig. 4f). Ontology analysis of the enriched proteins supported our hypothesis, with significant compartmental enrichment of the nucleus and terms associated with RNA binding, processing and splicing (Fig. 4g). Pleasingly, some of our most enriched proteins include TIAL1, CAPRIN1, FUBP3, BANF1, and PABPC1 (Fig. 4f) which have been previously implicated in the antiviral response^24,25^. We also noted enrichment of CASC3, EIF4A3, and RBM8A, which are crucial components of the exon junction complex. We validated CASC3 movement in response to dsRNA by immunofluorescence microscopy, demonstrating a shift from diffuse cytoplasmic localization to an enrichment in nuclear speckles, suggesting a potential role for the EJC in mediating the response to viral infection^26^ (Fig. 4h). We confirmed the enrichment of other hits in the nucleus in response to infection, including the known RBPs PABPC1 and EIF4A1, and the transcription factor JUNB, suggesting potential transcriptional regulation by the AP-1 complex.

### Interrogating the RUNX3 cistrome using FlAsH PL

The transcription factor RUNX3 is a 44 kDa protein that is required for development of tyrosine kinase receptor C proprioceptive neurons in dorsal root ganglia^27^, the specification of mature CD8^+^ T lymphocytes in the thymus^28,29^, and the differentiation of naive CD8^+^ T lymphocytes into long-lived memory cells following activation in response to infections^30,31^. RUNX3 DNA-binding requires physical cooperation through its RUNT-domain with a heterodimeric partner CBFB and its transcriptional activity is mediated by several regulatory domains that appear to recruit distinct co-regulatory factors^32^. However, the co-regulators with which RUNX3 associates in cells to mediate its chromatin-level regulation are ill-defined. Endogenous RUNX3 is strongly associated with chromatin and the nuclear matrix and is relatively insoluble under standard conditions to extract nuclear proteins. This characteristic is a barrier to immune-purification of RUNX3 from nuclear lysates and identification of associated factors, which might only weakly or transiently interact with RUNX3. We reasoned a proximity labeling approach in live cells might overcome this limitation. Furthermore, we anticipated appending an enzymatic labeling module to RUNX3 might obscure some of its functions whereas the minimally invasive FT modification would be less detrimental and might be advantageous.

To examine these possibilities, RUNX3-WT, RUNX3-FT and RUNX3-TurboID (RUNX3-TID) constructs were cloned, expressed in HEK293T cells and assessed for their localization and capacity to mediate proximity labeling. All RUNX3 constructs showed nuclear localization and biotinylation by confocal microscopy (Supplementary Fig. 5a). Notably, however, the RUNX3-TID construct showed significant biotinylation prior to addition of exogenous biotin. MS analysis after enrichment of biotinylated proteins indicated substantial similarities and critical differences between RUNX3-FT and RUNX3-TID interactomes (Supplementary Fig. 5b). Both RUNX3-FT and RUNX3-TID identified

RUNX3, transducin-like Enhancer-of-split (TLE) proteins, and CBFB as the top enriched hits in both datasets, consistent with previous studies^29,33^ (Fig. 5a, c). This indicates that both methods can capture the core RUNX3 interactome. However, RUNX3-TID identified a total of 2155 enriched proteins, and only 52.5% of hits were localized in the nucleus. The ratio increases to 57.5% while selecting the top 40 enriched hits, ranking by composite score. In contrast, RUNX3-FT identified a limited number of candidate factors of which 28 out of 29 were nuclear, suggesting RUNX3-FT captures a more specific interactome at chromatin with significantly less noise.

**Fig 5.**
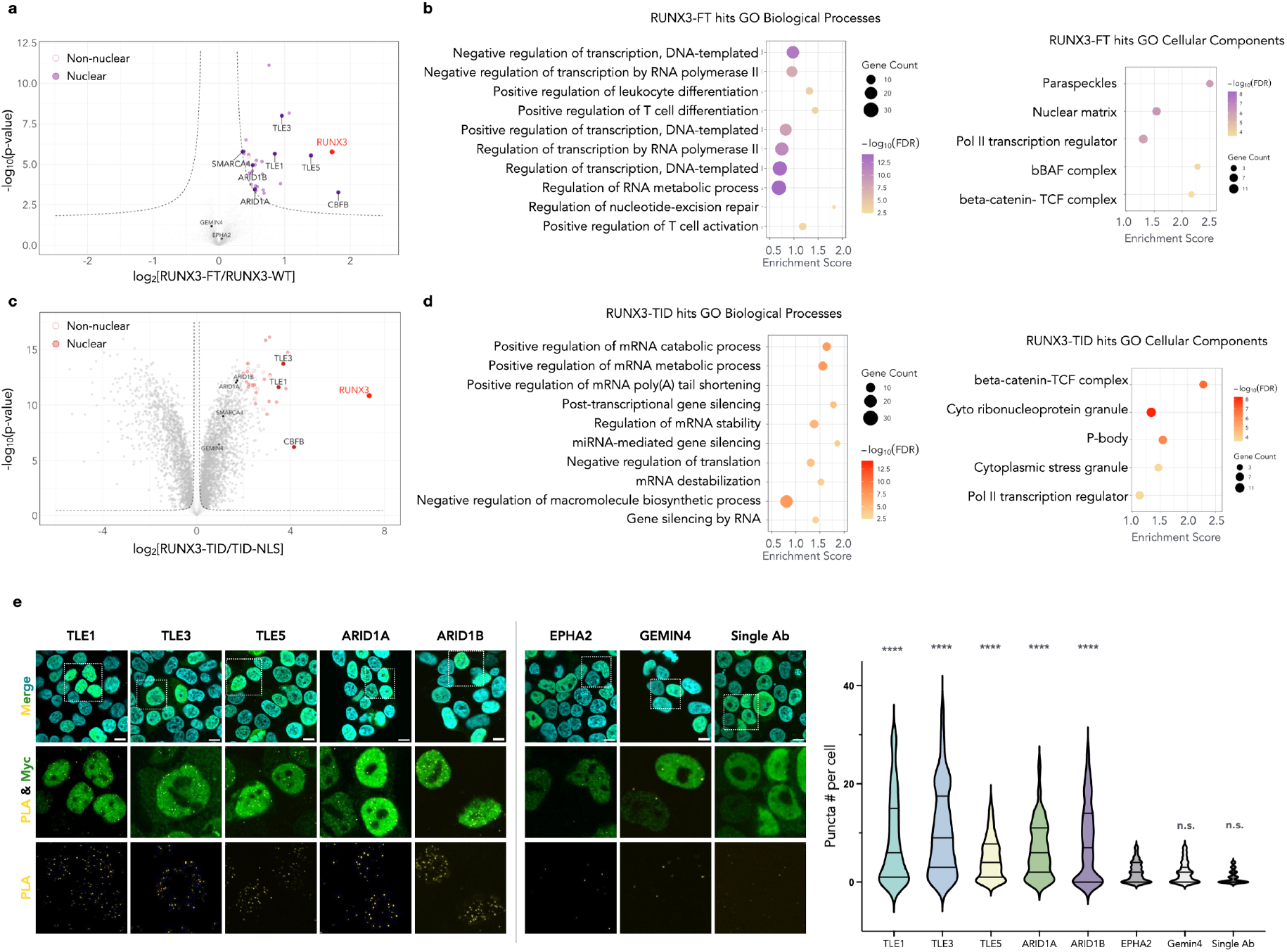
RUNX3 interactome is mapped by FlAsH-ID in HEK cells. **a**, A volcano plot derived from a two-sided *t*-test showing RUNX3 interactors from the FlAsH-ID method comparing to HEK cells overexpressing RUNX3-WT. Nuclear proteins, known interactors, and other components of the SWI/SNF complex and TLE protein family are highlighted. FDR<0.05. **b**, GO biological processes and cellular components analysis. Terms are ranked by a weighted harmonic mean between the enrichment score and -log_10_FDR. **c**, A volcano plot derived from a two-sided *t*-test showing RUNX3 interactors from the TurboID method comparing to HEK cells overexpressing TID-NLS. Hits with the top 40 composite score (log_2_(FC) * -log_10_(p-value)) were highlighted for direct comparison between datasets. Nuclear proteins, known interactors, and other components of the SWI/SNF complex and TLE protein family are highlighted. FDR<0.05. **d**, GO biological processes and cellular components analysis. Terms are ranked by a weighted harmonic mean between the enrichment score and -log_10_FDR. **e**, Representative images and violin plot with median and quartile lines comparing interaction between hits and RUNX3. EPHA2 and GEMIN4 were detected in the FlAsH dataset but not enriched. Scale bar = 10 µm (n>140, one-way ANOVA test followed by post hoc Dunnett’s multiple comparisons comparing to EPHA2 control. \*\*\*\**P*<0.0001).

GO enrichment analysis indicated that top candidates from each method were enriched with distinct molecular ontologies. Genes related to transcriptional regulation and T-cell differentiation were enriched in the FlAsH-ID dataset (Fig. 5b). Multiple key factors (ARID1A, ARID1B, and BRG1) from the SWI/SNF BRG1-associated factor (BAF) family of chromatin remodelers were specifically enriched using RUNX3-FT. While a functional relationship has been revealed between ARID1A and RUNX3^31^, no direct physical interaction has been reported. Comparison of ranked gene lists from TurboID and FlAsH-ID experiments showed that while many of the hits enriched in FlAsH-ID were also captured by TurboID analysis, they were not among the top enriched hits, e.g., BRG1 is ranked 25^th^ in FlAsH-ID but 341^st^ in TurboID (Supplementary Fig. 5c). Furthermore, the GO analysis of top TurboID hits showed enrichment of complexes and terms associated with RNA processing, ribonuclear protein complexes and cytoplasmic stress granules which are not known functions of RUNX3 (Fig. 5d). To account for RUNX3 regulation at the transcriptional level, we ran second TurboID labeling experiment with RUNX3-WT as the control. While RUNX3 and TLE proteins were again among the more enriched proteins, many of the hits enriched by FlAsH-ID remained buried in the gene list or not enriched at all. In addition, GO analysis indicates that most hits were relevant to RNA processing rather than transcriptional regulation (Supplementary Fig. 5d, e, f). The dramatic differences in interactome produced by these two systems demonstrates the capacity of our approach for interrogation of small transcriptionally active proteins such as RUNX3 (Supplementary Fig. 5g).

To demonstrate the veracity of the RUNX3 interactome provided by FlAsH-ID, we validated a selection of the most enriched proteins using proximity ligation assays. Interactions between RUNX3 and TLE1, 3, 5 in addition to ARID1A and ARID1B were all validated using this approach, demonstrating significant increases in signal when compared to single antibody controls and negative control proteins EPHA2 and GEMIN4 (Fig. 5e). While both EPHA2 and GEMIN4 show no enrichment in the FlAsH-ID analysis, TurboID labeling identified GEMIN4 as a significant hit (Fig. 5a, c).

The distinct proximity labelling characteristics of RUNX3-FT and RUNX3-TID suggested that each modified protein functioned differently in cells. To determine whether RUNX3-FT or RUNX3-TID altered the natural function of RUNX3, T cell receptor stimulated naive CD8^+^ T cells were transduced with retroviruses expressing unmodified wildtype RUNX3 (RUNX3-WT), RUNX3 with FT tag insertion (RUNX3-FT) or with TurboID fused (RUNX3-TID) and cultured under conditions that promote distinct effector and memory cell-like states (Fig. 6a, Supplementary Fig. 6a). The expression of 10 differentiation markers was examined on transduced cells by flow cytometry (Supplementary Fig. 6b). As expected from previous analyses of RUNX3 in cultured CD8^+^ T cells^34^, cells cultured in low concentrations of IL-2 more highly expressed CD103 (*Itgae*), PD-1 (*Pdcd1*), LAG3 (*Lag3*) and CD69 (*Cd69*) and concomitantly less highly expressed CD127 (*Il7r*), SLAMF6 (*Slamf6*) and CD62L (*Sell*) when transduced with RUNX3-WT as compared to empty vector (EV)-transduced or untransduced cells (Fig. 6b, Supplementary Fig. 6b). Notably, cells transduced with RUNX3-FT demonstrated a similar pattern of expression in these markers as RUNX3-WT transduced cells when cells were cultured in either low or high concentrations of IL-2, whereas cells transduced with RUNX3-TID manifested a distinct pattern. In particular, RUNX3-TID hyperexpressed CD69 and LAG3, failed to downregulate SLAMF6, CD127 and CD62L and less strongly expressed PD-1 and TIM3. Unsupervised clustering indicated cells transduced with RUNX3-WT or RUNX3-FT exhibited analogous behavior whereas those transduced with RUNX3-TID were more analogous to EV-transduced or untransduced cells (Fig. 6b). Thus, the typical gain-of-function induced by enforced expression of RUNX3-WT is closely mimicked by enforced expression of RUNX3-FT but not RUNX3-TID.

**Fig 6.**
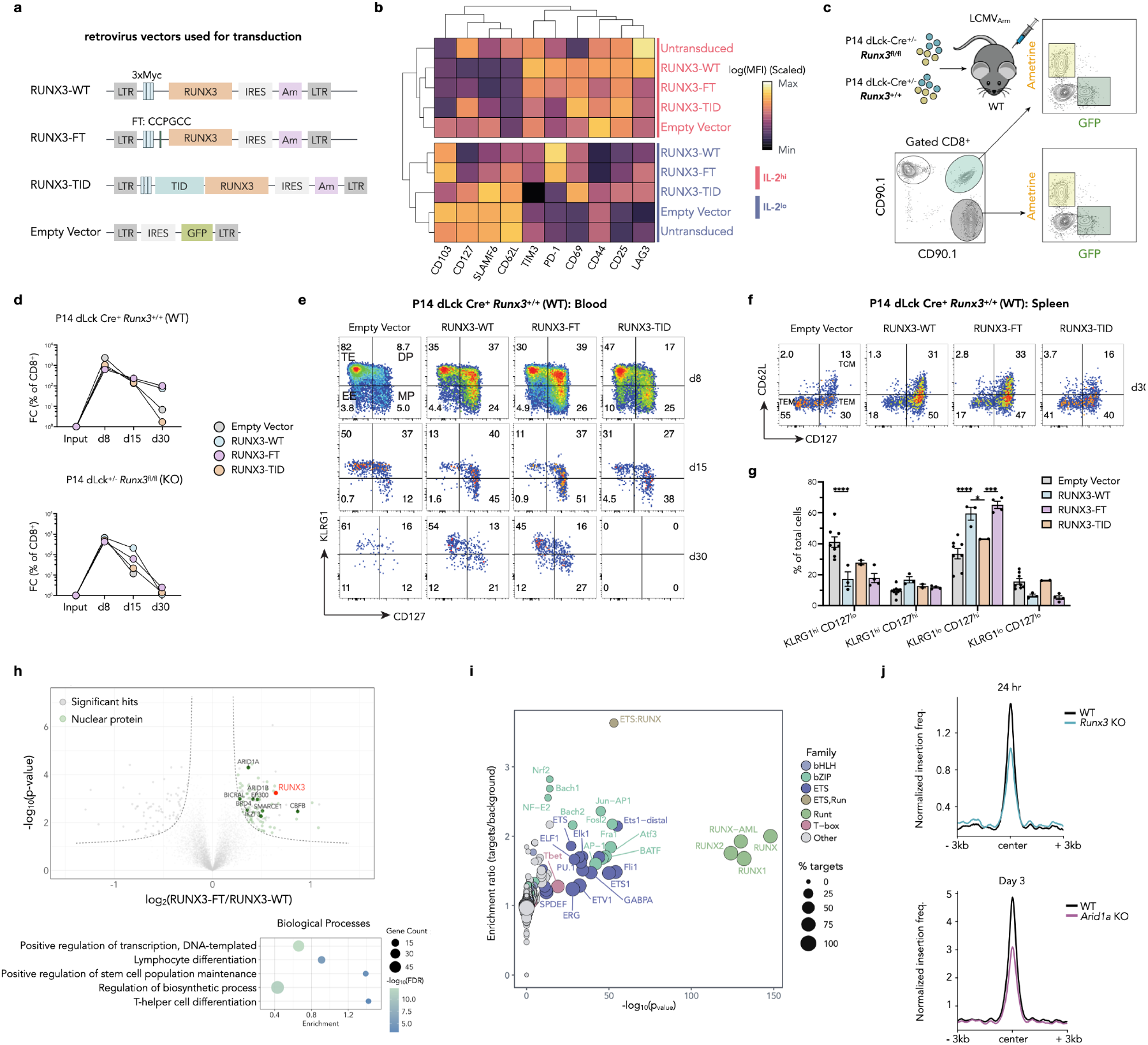
FlAsH-ID conserves RUNX3 biology and identified ARID1A as its interactor in CD8^+^ cells. **a**, Schematic of retroviral vectors used to transduce murine CD8^+^ T cells. **b**, Heatmap of log(Mean fluorescence intensity) during high or low IL-2 culture. Value of log(MFI) is scaled by column. **c**, Schematic of LCMV_Arm_ infection co-transfer experiment. GFP (Empty Vector) or Ametrine (RUNX3-WT, RUNX3-FT, RUNX3-TID) vectors were transduced into naïve CD8 T cells purified from P14 dLck-Cre^+^ *Runx3*^fl/fl^ or *Runx3*^+/+^ mice. **d**, Median fold change of transferred P14 populations expressed as a percentage of total CD8^+^ T cells normalized with input. **e**, Flow cytometry plots show representative KLRG1/CD127 staining of gated P14 cells in the blood 8, 15, or 30 days post-infection (p.i.). **f**, Flow cytometry plots show representative CD127/CD62L staining of gated P14 cells to enumerate central memory (TCM) and effector memory (TEM) populations in the spleen 30 days p.i.. **g**, Frequencies of KLRG1/CD127 subsets in the spleen 30 days p.i. are summarized (two-way ANOVA test followed by Tukey’s multiple comparisons test for differences between conditions. Not significant comparisons are not shown. \**P*<0.05, \*\**P*<0.01, \*\*\**P*<0.001, \*\*\*\**P*<0.0001) **h**, Top, a volcano plot derived from a two-sided *t*-test showing RUNX3 interactors from the FlAsH-ID method comparing to CD8^+^ T cells overexpressing RUNX3-WT. Nuclear proteins, known interactors, and components of the SWI/SNF complex are highlighted. FDR<0.05. Bottom, GO biological processes analysis of enriched hits. Terms are ranked by a weighted harmonic mean between the enrichment score and -log_10_FDR. Terms with similarity ≥ 0.7 are merged. **i**, Enrichment ratio of motifs in activation inducible peaks occupied by both RUNX3 and ARID1A (normalized to background signal, HOMER motif analysis). Motif families are grouped by color. **j**, Normalized insertion frequency (ATAC) of *Runx3* KO and *Arid1a* KO CD8^+^ T cells during early response to LCMV_Arm_. Insertion frequency was normalized to the corresponding WT control within the same experiment.

In addition to altering the function of RUNX3, we anticipated that fusion of TurboID to RUNX3 might compromise its utility for analysis of cells *in vivo* because expression of foreign genes (e.g., selectable markers or gene editing machinery) in murine cells that are transferred to immunocompetent mice can lead to their rejection^35^. To determine whether RUNX3-FT might be suitable for analysis of CD8^+^ T cells during immune responses *in vivo*, congenically mismatched wildtype (Thy1.1^+^) and *Runx3*-deficient (Thy1.1^+^ 1.2^+^) TCR transgenic P14 CD8^+^ T cells which recognize the GP33 epitope from *Lymphocytic choriomeningitis virus* (LCMV) were complemented by retroviral transduction with EV, RUNX3-WT, RUNX3-FT, or RUNX3-TID. Wildtype and *Runx3*-deficient P14 cells transduced with RV-EV were mixed with those transduced with each RUNX3 variant and co-adoptively transferred to naive, congenic C57BL6/J mice (Thy1.2^+^) wildtype hosts and these recipients were subjected to infection with the Armstrong strain of LCMV (LCMV_Arm_) (Fig. 6c). In this setting, LCMV_Arm_ causes an infection that is acutely resolved by the wildtype host immune response independently of the adoptively transferred donor P14 cells. This approach allows determining the cell-intrinsic effects caused by the retrovirally transduced genes in control and perturbed P14 cells in the same hosts that resolve the infection normally. By day 8 post infection (p.i.), which is near when the maximum accumulation of CD8^+^ T cells occurs, wildtype P14 cells transduced with RUNX3-WT, RUNX3-FT and RUNX3-TID accumulated similarly to each other and slightly less than those transduced with EV (Fig. 6d, top panel), consistent with the slight negative effect of enforced RUNX3 activity on cell proliferation^30^. In addition, all three versions of RUNX3 were able to rescue the impaired accumulation of *Runx3*-deficient P14 cells as compared to those that were EV transduced. Therefore, RUNX3-FT and RUNX3-TID complemented defective accumulation of *Runx3*-deficient CD8^+^ T cells near the peak response. However, the accumulation of *Runx3*-deficient P14 cells transduced with RUNX3-TID was substantially reduced on days 15 and 30 p.i. compared to those transduced with either RUNX3-WT or RUNX3-FT, and was comparable to those transduced with EV. In contrast, *Runx3*-deficient P14 cells transduced with either RUNX3-WT or RUNX3-FT accumulated to a similar degree as wildtype P14 cells transduced with EV, suggesting they both fully rescued the accumulation defect that is characteristic of *Runx3* deficient CD8^+^ T cells (Fig. 6d, bottom panel). In addition, further phenotypic analysis demonstrated that RUNX3-WT and RUNX3-FT, but not RUNX3-TID, was able to increase the fraction of KLRG1^hi^ CD127^hi^ “double positive” effector cells, which absolutely require *Runx3* for development^30^, in wildtype P14 cells at all time points (Fig. 6e, Supplementary Fig. 6c). Analysis of splenic populations during the transition to the memory phase (30d p.i.) revealed that both RUNX3-WT and RUNX3-FT caused a fractional increase in CD62L^hi^ CD127^hi^ central memory (TCM) cells, and increased overall fractional abundance of CD62L^hi^ cells (Fig. 6f, Supplementary Fig. 6d) Further, transduction with RUNX3-TID failed to increase the fraction of KLG1^lo^CD127^hi^ memory cells (Fig. 6g). Thus, RUNX3-FT, but not RUNX3-TID, functions analogously to wildtype RUNX3 in the formation of effector cells that give rise to long-lived memory CD8^+^ T cells. These results suggest that RUNX3-FT conserves the biology of wildtype RUNX3 and is likely to be valuable for identifying molecules in primary cells *in vitro* and *in vivo*.

To extend the proximity labeling results from HEK cells and identify putative RUNX3 co-factors in primary CD8^+^ T cells we focused on RUNX3-FT. FlAsH-ID analysis of the RUNX3-FT transduced CD8^+^ T cells identified 110 proteins that were significantly enriched relative to RUNX3-WT cells (Fig. 6h). RUNX3 and CBFB were the two of the most enriched candidates, as expected. In addition, consistent with FLAsH-ID analysis of RUNX3 in HEK293T, members of the BAF complex, including ARID1A, ARID1B, SMARCE1, BICRAL, and DPF2 were enriched in proximity to RUNX3-FT in activated CD8^+^ T cells. Accordingly, GO terms related to SWI/SNF complexes, the nucleus and positive regulation of stem cell maintenance, transcription and T cell differentiation were all enriched within the RUNX3-FT interactome (Fig. 6h, Supplementary Fig. 6e). Also consistent with these results, BRD4 and EP300 whose functions in CD8^+^ T cells have been positively correlated with cell states that depend on RUNX3 for development were also enriched in conjunction with RUNX3^36^. These results suggest that expression of RUNX3-FT and FlAsH-ID are likely to have identified *bona fide* regulators that cooperate with RUNX3 in primary CD8^+^ T cells.

In naive CD8^+^ T cells, RUNX3 is required to promote chromatin accessibility at the majority of *cis*-regulatory regions that gain accessibility *de novo* during initial T cell receptor (TCR) stimulation^30^. A large proportion of these regions are stably accessible in mature memory CD8^+^ T cells suggesting RUNX3 initially establishes their accessibility. Analogously, ARID1A is also necessary to establish chromatin accessibility at a large number of *cis*-regulatory regions in TCR-stimulated CD8^+^ T cells during acute viral infection^34^. The enrichment of ARID1A in proximity to RUNX3 was consistent with these results and implied that both factors might promote chromatin remodeling during initial TCR stimulation of naive CD8^+^ T cells. Consistent with this hypothesis, genomic regions that become RUNX3 occupied in activated CD8^+^ T cells relative to unstimulated CD8^+^ T cells from naive mice overlapped extensively with regions that gain ARID1A occupancy in activated, but not naive, CD8^+^ T cells (Supplementary Fig 6f). Thus, more than half of genomic regions that bind RUNX3 after naive CD8^+^ T cells are initially activated also gain ARID1A-occupancy following activation. In line with this observation, DNA sequences recognized by RUNX-TFs were the most highly enriched TF motifs at these co-occupied regions (Fig 6i). To determine whether chromatin accessibility that is induced at these regions required either RUNX3 or ARID1A, the ATAC-seq signals at these co-occupied regions were examined in *Runx3*-deficient and *Arid1a*-deficient CD8^+^ T cells compared to their paired wildtype control CD8^+^ T cells following TCR stimulation (Fig 6i). CD8^+^ T cells lacking either RUNX3 or ARID1A each exhibited strongly impaired chromatin accessibility at these regions, indicating that each factor is necessary for promoting chromatin accessibility of these regions during initial TCR stimulation. Notably, many of these regions comprised *cis*-regulatory sequences in genes that are essential for differentiation of activated CD8^+^ T cells, such as *Il2ra* (Supplementary Fig 6f, g). These results indicate RUNX3 and ARID1A concertedly drive chromatin remodeling in activated CD8^+^ T cells in *cis*-regulatory regions that promote their differentiation into protective effector and memory CD8^+^ T cells.

## Discussion

Proximity labeling has become a mainstay technique in molecular biology, elucidating the interactomes of proteins and nucleic acids in countless contexts. Although dozens of techniques are now available for live cell proximity labeling, all methods rely on the use of large fusion proteins that have the potential to perturb native function. Our approach, FlAsH-ID, provides equivalent data to best-in-class PL methods but, through the repurposing of the Tsien FlAsH imaging technology, requires the insertion of only a 6 amino acid tag. We demonstrate that this approach is broadly applicable to different cellular compartments and can generate accurate interactomes in model cell lines. Furthermore, once incorporated into live cells, the labeled proteins are stable for at least 24 hours, enabling time resolved, stimuli responsive proximity labeling experiments.

We demonstrate that in contrast to conventional PL enzymes, FlAsH-ID does not disrupt the phenotype of the transcription factor RUNX3 in primary T cells and in vivo. Furthermore, PL of RUNX3 in primary CD8+ T cells identified a direct interaction and functional relationship with BAF complex in remodeling chromatin.

These data position FlAsH-ID as a valuable technology that can be readily employed across the chemical and molecular biology communities to study myriad sensitive proteins that are currently recalcitrant with existing interactomics approaches. Furthermore, in contrast to many other photoproximity labeling methods, all components required for use are commercially available, democratizing the use of this method to users without synthetic capability. Future work will focus on expanding the available catalysis modes to provide variable radii proximity labeling and exploring the generality of this platform across different cellular and tissue contexts.

## Supporting information

Table 1: Plasmid sequences

Table 2: Complied Processed Mass spec file

Table 3: Protein groups file

Table 4: Peptide groups file

## Supplementary Information

Supplementary information is linked to the online version of the paper.

## Data Availability

All relevant data are included in the manuscript and supplementary information. Mass spectrometry data files have been uploaded to the MassIVE proteomics database (MSV000099500).

## Acknowledgements

Research reported in this publication was supported by the Office of The Director, of the National Institutes of Health under Award Number S10OD036363, the National Institute of General Medical Sciences of the National Institutes of Health (R35GM150765) and the National Institute of Allergy and Infectious Diseases (P01AI145815). The content is solely the responsibility of the authors and does not necessarily represent the official views of the National Institutes of Health. CPS also acknowledges the Wertheim UF-Scripps for start-up funds. The authors thank George Tsaprailis and Catherina Scharager Tapia at the UF-Scripps Proteomics Facility, Robert M. Witwicki, Li Pan, and Marlene L. Biller at the UF-Scripps Genomics Facility, and Gogce C. Cryen and Alexander Trouern-Trend at the Bioinformatics and Statistics Core.

## Author Contributions

CPS and MEP conceived the work. CPS, MEP, WZ, SN, and JK, designed and executed the experiments and TV analyzed data. CPS, WZ, SN, and MEP prepared the manuscript.

## Author Information

The authors declare no competing financial interests. Readers are welcome to comment on the online version of the paper. Correspondence and requests for materials should be addressed to CPS or MEP.

## References

1) Qin, W., Cho, K. F., Cavanagh, P. E. & Ting, A. Y. Deciphering molecular interactions by proximity labeling. Nat. Methods 18, 133–143 (2021).

2) Milione, R. R., Schell, B.-B., Douglas, C. J. & Seath, C. P. Creative approaches using proximity labeling to gain new biological insights. Trends Biochem. Sci. 49, 224–235 (2024).

3) Seath, C. P. et al. Tracking chromatin state changes using nanoscale photoproximity labelling. Nature 616, 574–580 (2023).

4) Lin, Z. et al. Multiscale photocatalytic proximity labeling reveals cell surface neighbors on and between cells. Science 385, eadl5763 (2024).

5) Zhou, Y. & Zou, P. The evolving capabilities of enzyme-mediated proximity labeling. Curr. Opin. Chem. Biol. 60, 30–38 (2021).

6) Kim, D. I. et al. An improved smaller biotin ligase for BioID proximity labeling. Mol. Biol. Cell 27, 1188–1196 (2016).

7) Fatti, E., Khawaja, S. & Weis, K. The dark side of fluorescent protein tagging-the impact of protein tags on biomolecular condensation. Mol. Biol. Cell 36, br10 (2025).

8) Pandey, N. K. et al. Fluorescent protein tagging promotes phase separation and alters the aggregation pathway of huntingtin exon-1. J. Biol. Chem. 300, 105585 (2024).

9) Knutson, S. D., Buksh, B. F., Huth, S. W., Morgan, D. C. & MacMillan, D. W. C. Current advances in photocatalytic proximity labeling. Cell Chem. Biol. 31, 1145–1161 (2024).

10) Tong, F., Zhou, W., Janiszewska, M. & Seath, C. P. Multiprobe Photoproximity Labeling of the EGFR Interactome in Glioblastoma Using Red-Light. J. Am. Chem. Soc. 147, 9316–9327 (2025).

11) Buksh, B. F. et al. μMap-Red: Proximity Labeling by Red Light Photocatalysis. J. Am. Chem. Soc. 144, 6154–6162 (2022).

12) Knutson, S. D. et al. Parallel proteomic and transcriptomic microenvironment mapping (μmap) of nuclear condensates in living cells. J. Am. Chem. Soc. 147, 488–497 (2025).

13) Griffin, B. A., Adams, S. R. & Tsien, R. Y. Specific covalent labeling of recombinant protein molecules inside live cells. Science 281, 269–272 (1998).

14) Hoffmann, C. et al. Fluorescent labeling of tetracysteine-tagged proteins in intact cells. Nat. Protoc. 5, 1666–1677 (2010).

15) Andresen, M., Schmitz-Salue, R. & Jakobs, S. Short tetracysteine tags to betatubulin demonstrate the significance of small labels for live cell imaging. Mol. Biol. Cell 15, 5616–5622 (2004).

16) Liu, B., Archer, C. T., Burdine, L., Gillette, T. G. & Kodadek, T. Label transfer chemistry for the characterization of protein-protein interactions. J. Am. Chem. Soc. 129, 12348–12349 (2007).

17) Taguchi, Y. et al. Specific biarsenical labeling of cell surface proteins allows fluorescent- and biotin-tagging of amyloid precursor protein and prion proteins. Mol. Biol. Cell 20, 233–244 (2009).

18) Lelek, M. et al. Superresolution imaging of HIV in infected cells with FlAsH-PALM. Proc Natl Acad Sci USA 109, 8564–8569 (2012).

19) Martin, B. R., Giepmans, B. N. G., Adams, S. R. & Tsien, R. Y. Mammalian cell-based optimization of the biarsenical-binding tetracysteine motif for improved fluorescence and affinity. Nat. Biotechnol. 23, 1308–1314 (2005).

20) Oughtred, R. et al. The BioGRID database: A comprehensive biomedical resource of curated protein, genetic, and chemical interactions. Protein Sci. 30, 187–200 (2021).

21) Rath, S. et al. MitoCarta3.0: an updated mitochondrial proteome now with suborganelle localization and pathway annotations. Nucleic Acids Res. 49, D1541– D1547 (2021).

22) Nakielny, S. & Dreyfuss, G. Transport of proteins and RNAs in and out of the nucleus. Cell 99, 677–690 (1999).

23) Richner, J. M. et al. Global mRNA degradation during lytic gammaherpesvirus infection contributes to establishment of viral latency. PLoS Pathog. 7, e1002150 (2011).

24) Burke, J. M. et al. RNase L activation in the cytoplasm induces aberrant processing of mRNAs in the nucleus. PLoS Pathog. 18, e1010930 (2022).

25) Gilbertson, S., Federspiel, J. D., Hartenian, E., Cristea, I. M. & Glaunsinger, B. Changes in mRNA abundance drive shuttling of RNA binding proteins, linking cytoplasmic RNA degradation to transcription. eLife 7, (2018).

26) Schlautmann, L. P. & Gehring, N. H. A day in the life of the exon junction complex. Biomolecules 10, (2020).

27) Levanon, D. et al. The Runx3 transcription factor regulates development and survival of TrkC dorsal root ganglia neurons. EMBO J. 21, 3454–3463 (2002).

28) Taniuchi, I. et al. Differential requirements for Runx proteins in CD4 repression and epigenetic silencing during T lymphocyte development. Cell 111, 621–633 (2002).

29) Woolf, E. et al. Runx3 and Runx1 are required for CD8 T cell development during thymopoiesis. Proc Natl Acad Sci USA 100, 7731–7736 (2003).

30) Wang, D. et al. The Transcription Factor Runx3 Establishes Chromatin Accessibility of cis-Regulatory Landscapes that Drive Memory Cytotoxic T Lymphocyte Formation. Immunity 48, 659-674.e6 (2018).

31) Milner, J. J. et al. Runx3 programs CD8+ T cell residency in non-lymphoid tissues and tumours. Nature 552, 253–257 (2017).

32) Mevel, R., Draper, J. E., Lie-A-Ling, M., Kouskoff, V. & Lacaud, G. RUNX transcription factors: orchestrators of development. Development 146, (2019).

33) Yarmus, M. et al. Groucho/transducin-like Enhancer-of-split (TLE)-dependent and -independent transcriptional regulation by Runx3. Proc Natl Acad Sci USA 103, 7384–7389 (2006).

34) Adelman, K. & Lis, J. T. Promoter-proximal pausing of RNA polymerase II: emerging roles in metazoans. Nat. Rev. Genet. 13, 720–731 (2012).

35) Chen, R. et al. In vivo RNA interference screens identify regulators of antiviral CD4(+) and CD8(+) T cell differentiation. Immunity 41, 325–338 (2014).

36) Milner, J. J. et al. Bromodomain protein BRD4 directs and sustains CD8 T cell differentiation during infection. J. Exp. Med. 218, (2021).

