## Supplementary Figures and data for "FlAsH-ID: A short peptide tag for live cell photoproximity labeling"

### Supporting Information

**Wuyue Zhou,[1,2] Shashank Nagaraja, [2,3] Jihye Kim[3], Thomas Venables[3], Matthew E. Pipkin,*,[2,3] Ciaran P. Seath*,[1,2]**

*[1]Department of Chemistry, Wertheim UF Scripps, Jupiter, Florida 33418, USA. [2]Skaggs Graduate School, The Scripps Research Institute, Jupiter Florida, 33418, USA. [3]Department of Immunology and Microbiology, Wertheim UF Scripps, Jupiter, Florida 33418, USA.*

#### Supplementary Figures

### Supporting Figure 1 – In vitro development of FlAsH-ID

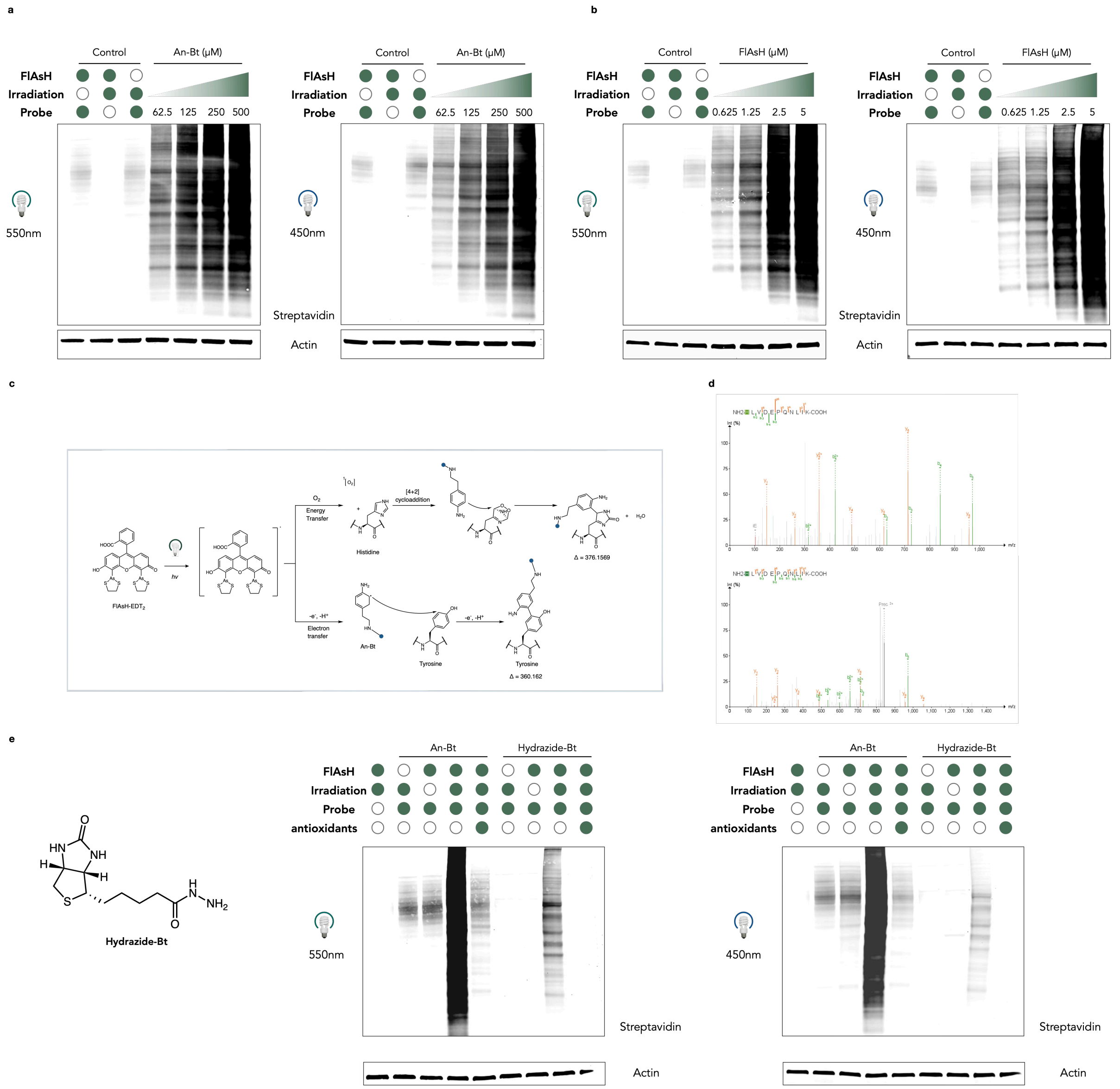

**Supporting Figure 1**. **a**, Activation of An-Bt with FlAsH-EDT$_2$ is dependent upon probe concentration. **b**, Activation of An-Bt with FlAsH-EDT$_2$ is dependent upon catalyst concentration. **c**, Potential mechanisms of FlAsH-EDT$_2$ activation. Following light administration, generated singlet oxygen locally oxidizes histidine residues, which forms endoperoxide intermediates that are trapped by the nucleophilic aniline probe and causes a mass shift of 376.16 Da on the histidine residue. Potentially, a photoexcited FlAsH could oxidize An-Bt through single electron transfer. This would reveal an aminyl radical that can intercept SOMOphilic amino acids like tyrosine and tryptophan and leads to a mass shift of 360.162 Da. **d**, Other MS2 spectra for peptide HLVDEPQNLIK showing modification of histidine 402 with the desired mass shift. **e,** FlAsH-

$EDT_2$ activates hydrazide-bt, and the signal could be quenched by addition of antioxidants (10mM sodium ascorbate, 5mM Trolox).

#### Supporting Figure 2 – Confirmation of FlAsH-ID with immunofluorescence microscopy

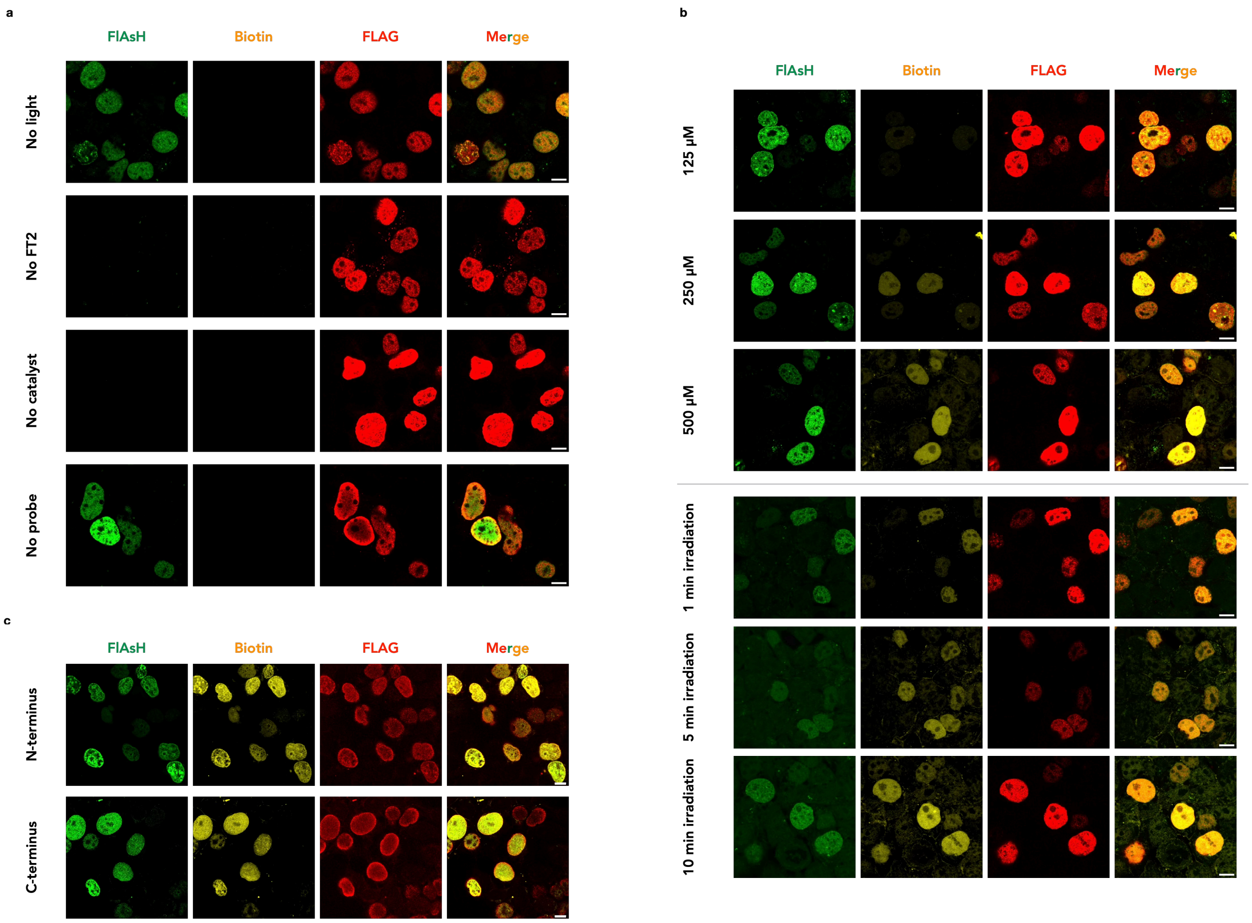

**Supporting Figure 2**. **a**, Biotinylation signal is only observed when all components were present. **b**, Biotinylation signal increases with longer irradiation and higher probe concentration. **c**, FlAsH tag (FT) incorporation is tolerated at both termini. Scale bar = 10 µm.

### Supporting Figure 3 – FlAsH-ID labeling of H3.1 in HEK cells

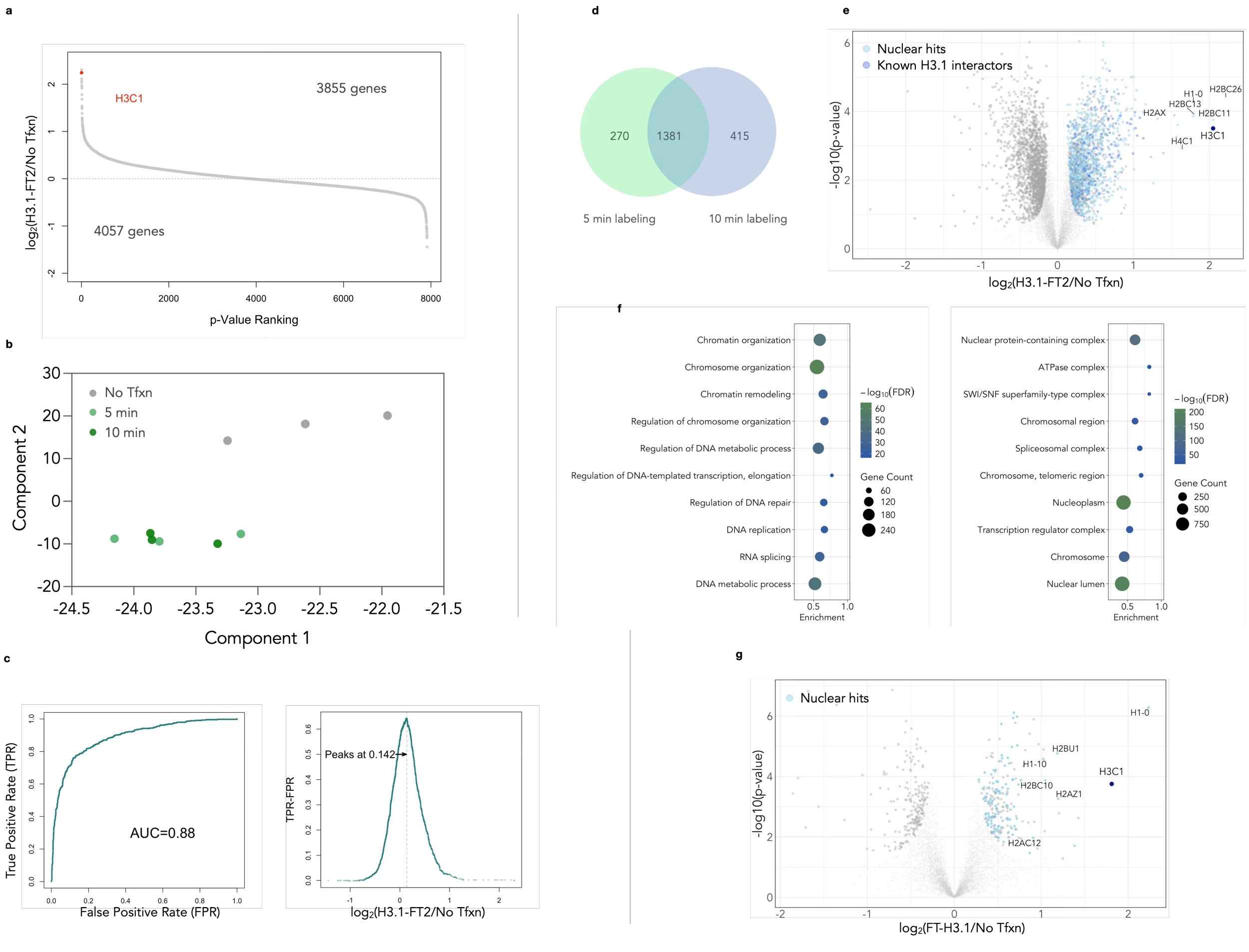

**Supporting Figure 3**. **a**, Waterfall plot of H3.1 FlAsH-ID dataset. H3.1 was among one of the highest enriched hits. **b**, Principal component analysis of the experiment. **c**, ROC analysis and cut-off determination. **d**, Overlapping hits between 5 min and 10 min treatment. **e**, A volcano plot derived from a two-sided *t*-test showing H3.1 interactors from FlAsH-ID while labeling for 10 min. FDR<0.05. Components of nucleosome were labeled on the plot. **f**, GO biological processes analysis for 10 min. **g**, A volcano plot derived from a two-sided *t*-test showing H3.1 interactors from FlAsH-ID while FT incorporated at the N-terminus histone tail. Components of nucleosome were labeled on the plot.

### Supporting Figure 4 – U-2 OS proteomic workflow and sample validation

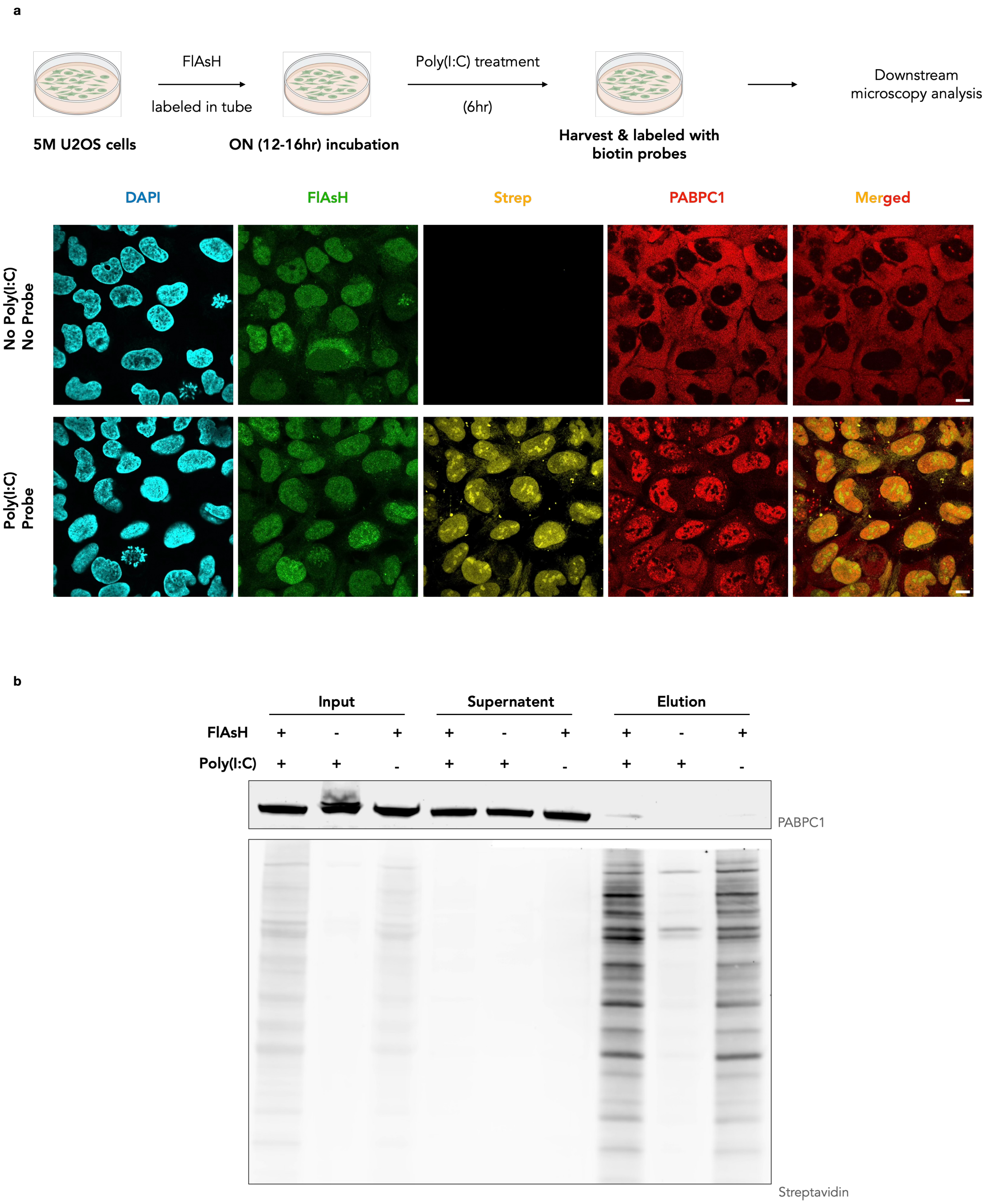

**Supporting Figure 4**. **a**, General workflow of the U-2 OS FlAsH-ID labeling experiment. Labeling was confirmed by biotinylation signal, and effect of poly(I:C) confirmed by PABPC1 translocation. **b**, Western blot confirmation of the proteomic sample. 5% of the eluted protein were run for WB analysis.

#### Supporting Figure 5 – RUNX3 FlAsH-ID in HEK293T cells

**a**

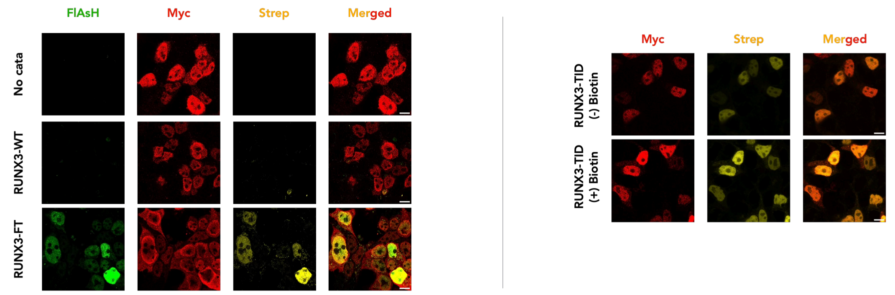

**b**

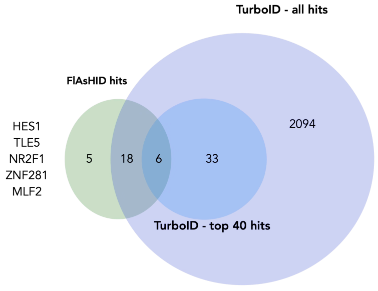

**c**

| | FlAsH-ID | TurboID |
|---|---|---|
| TLE4 | 3 | 13 |
| TLE3 | 5 | 3 |
| CBFB | 6 | 34 |
| TLE1 | 7 | 7 |
| ATN1 | 8 | 12 |
| PSPC1 | 9 | 333 |
| SSBP3 | 11 | 79 |
| NONO | 12 | 215 |
| RFX1 | 13 | 142 |
| ARID1B | 14 | 77 |
| TCF20 | 15 | 1112 |
| PIAS1 | 16 | 376 |
| NR2F2 | 19 | 1946 |
| FOXC1 | 20 | 1457 |
| CREB1 | 21 | 127 |
| HNRNPM | 22 | 337 |
| ZBTB10 | 23 | 1366 |
| ZIC2 | 24 | 1570 |
| SMARCA4 | 25 | 341 |
| SFPQ | 26 | 363 |
| DLX5 | 27 | 1555 |
| ARID1A | 28 | 91 |
| HSPA6 | 29 | 6 |
| ZNF609 | 30 | 324 |

**d**

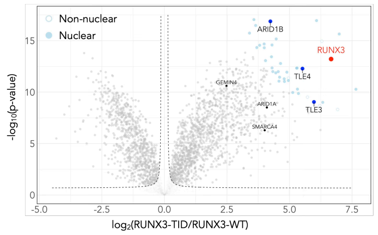

**e** **f** **g**

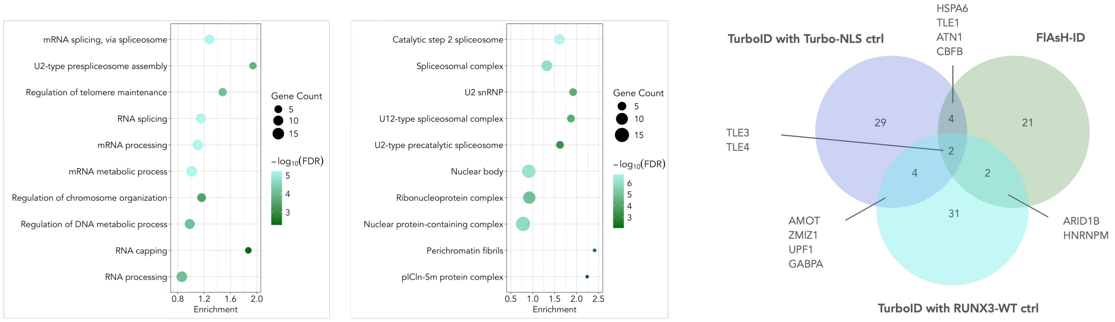

**Supporting Figure 5**. **a**, Immunofluorescence (IF) confirmation of FlAsH-ID and TurboID with RUNX3 in HEK cells. Background labeling is observed for TurboID without biotin supplement. **b,** Overlap between the TurboID hits and the FlAsH-ID hits. For TurboID datasets, hits with the top 40 composite score ($\log_2$(FC) * $-\log_{10}$(p-value)) were selected and labeled on the plot. **c**, Rankings of overlapping hits between TurboID and FlAsH-ID. **d**, A volcano plot derived from a two-sided *t*-test showing RUNX3 interactors from Turbo-ID while cells overexpressing RUNX3-WT were used as controls. FDR<0.05. **e**, GO biological processes. Terms are ranked by a weighted harmonic mean between the enrichment score and $-\log_{10}$FDR. **f**, GO cellular components. Terms are ranked by a weighted harmonic mean between the enrichment score and $-\log_{10}$FDR. **g**, Overlap across three datasets.

#### Supporting Figure 6 – RUNX3 FlAsH-ID in CD8$^+$ T cells

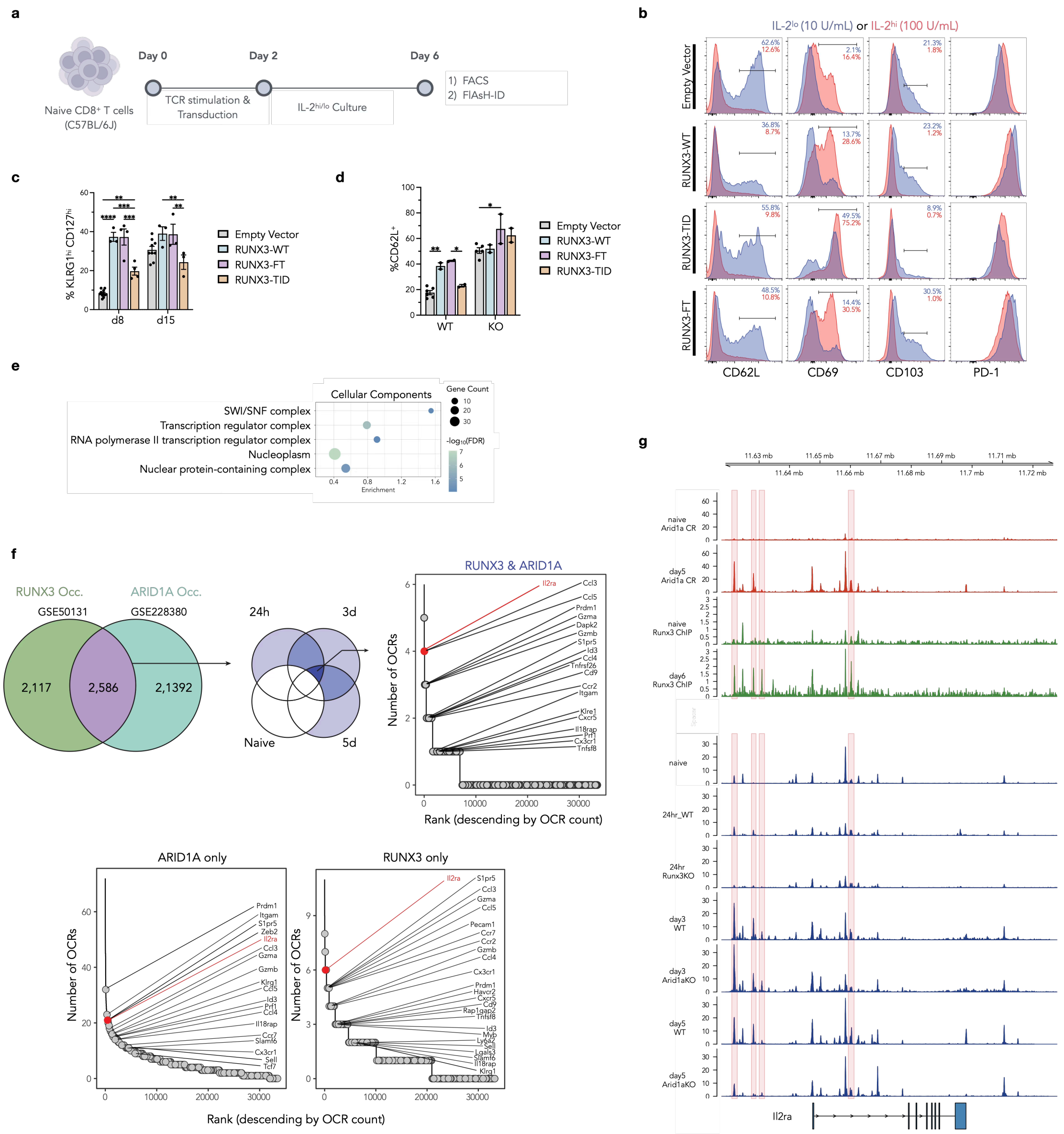

**Supplementary Figure 6. a**, Schematic for *ex vivo* stimulation and transduction of CD8$^+$ T cells. **b**, Fluorescence intensity histograms of T cell markers following plate-bound anti-CD3/CD28 stimulation. **c**, Percentage frequency of KLRG1$^{hi}$CD127$^{hi}$ double positive (DP) effector cells in the 4 transduced populations (P14 *Runx3*$^{+/+}$ cells) (two-way ANOVA test followed by Tukey's multiple comparisons test for differences between conditions. Not significant comparisons are not shown. **P*<0.05, ***P*<0.01, ****P*<0.001, *****P*<0.0001). **d**, Frequency of CD62L$^{hi}$ cells at 30d post-infection (splenocytes) (two-way ANOVA test followed by Tukey's multiple comparisons test for differences between conditions). **e**, GO cellular components analysis of enriched hits. Terms are ranked by a weighted harmonic mean between the enrichment score and -log$_{10}$FDR. Terms with similarity ≥ 0.7 are merged. **f**, Top: filtering schematic of TCR stimulation-inducible peaks, followed by designation of RUNX3 only/ARID1A only/both/none occupancy (RUNX3: ChiP, ARID1A: CUT&RUN). Inducible ATACseq regions were defined as those with log$_2$(FC) >1 and FDR < 0.05 at 24hr, 3d, or 5d relative to naïve. Bottom: ranked distribution of Open Chromatin Regions (OCRs) assigned per gene for both RUNX3 & ARID1A occupied, RUNX3 only occupied, and ARID1A only occupied peaksets, **g**, Genomic tracks of ARID1A (CUT&RUN) (naïve, d5 $LCMV_{Arm}$), Runx3 ChIP (naïve, d5 $LCMV_{Arm}$), ATAC (naïve, WT/*Runx3*KO 24h ex vivo TCR stim), ATAC (WT/*Arid1a*KO at 3, 5d $LCMV_{Arm}$).

#### General Considerations

Water was purified using a Millipore Milli-Q Integral Water Purification System. All buffers and synthetic starting materials were used as received from commercial sources. Eppendorf Protein LoBind tubes (Z666505) were purchased from Millipore Sigma. RIPA Lysis buffer (10X, 20-188) was purchased from Millipore Sigma. Halt™ Protease Inhibitor Cocktail (100X, 78438), Pierce BCA Protein Assay Kit (23227), Bolt Bis-Tris Plus Gel (NW04120BOX) and iBright Prestained Protein ladder (LC5615) were purchased from Thermo Fisher Scientific. TBST (20X, TBST01-03) was purchased from Bioland Scientific LLC. DMEM (Gibco, 10566016), DPBS (Gibco, 14190250), Fetal Bovine Serum (Gibco, 10437-028) and Trypsin-EDTA (Gibco, 25300054) were obtained from Thermo Fisher Scientific. Trypsin Protease (MS grade, 90057) was purchased from Thermo Fisher Scientific. Streptavidin Mag Sepharose magnetic beads (28985799) were purchased from Cytiva. Duolink® Proximity Ligation Assay kits (Duolink® In Situ PLA® Probe Anti-Rabbit PLUS, DUO92002; Duolink® In Situ PLA® Probe Anti-Mouse MINUS, DUO92004; Duolink® In Situ Detection Reagents Orange, DUO92007) and Acetonitrile (271004) were purchased from Millipore Sigma. FlAsH-$EDT_2$ (20704) was purchased from Cayman Chemical Co Inc.

#### Antibodies used in this study.

Western blot

| | | |
|---|---|---|
| Anti-FLAG | Cell Signaling (8146S) | 1:1,000 in TBST |
| Anti-Myc | Life Technologies (A190104A) | 1:1,000 in TBST |
| Anti-HDAC1 | Cell Signaling (34589T) | 1:1,000 in TBST |
| Anti-actin | Abcam (ab31830) | 1:5,000 in TBST |
| Anti-PABPC | Fisher Scientific (NC1999506) | 1:2000 in TBST |
| Anti-Mouse | LI-COR IRDye 680 nm | 1:10,000 in TBST |
| Anti-Rabbit | LI-COR IRDye 488 nm | 1:10,000 in TBST |
| Streptavidin | LI-COR IRDye Strep-800 nm | 1:10,000 in TBST |

Immunofluorescence

| | | |
|---|---|---|
| Anti-FLAG | Cell Signaling (8146S) | 1:500 in 2% BSA in PBST |
| Anti-Myc | Life Technologies (A190104A) | 1:1000 in 2% BSA in PBST |
| Anti-PABPC | Fisher Scientific (NC1999506) | 1:2,000 in 2% BSA in PBST |
| Anti-CASC3 | Life Technologies (A302472AT) | 1:500 in 2% BSA in PBST |
| Anti-EIF4A1 | Life Technologies (MA551507) | 1:500 in 2% BSA in PBST |
| Anti-JUNB | Cell Signaling (3573S) | 1:500 in 2% BSA in PBST |
| Anti-Mouse | LI-COR IRDye 488 nm | 1:1000 in 2% BSA in PBST |
| Anti-Rabbit | LI-COR IRDye 488 nm | 1:1000 in 2% BSA in PBST |
| Anti-Mouse | LI-COR IRDye 647 nm | 1:1000 in 2% BSA in PBST |
| Anti-Rabbit | LI-COR IRDye 647 nm | 1:1000 in 2% BSA in PBST |
| Streptavidin | LI-COR IRDye Strep-550 nm | 1:1000 in 2% BSA in PBST |

PLA

| | | |
|---|---|---|
| Mt Anti-Myc | Life Technologies (A190104A) | 1:500 in antibody diluent |
| Gt Anti-Myc | Life Technologies (A190104A) | 1:200 in antibody diluent |
| Anti-ARID1A | Life Technologies (PA585568) | 1:400 in antibody diluent |
| Anti-ARID1B | Fisher Scientific (NC2457300) | 1:1000 in antibody diluent |
| Anti-TLE1 | Life Technologies (UM870143) | 1:100 in antibody diluent |
| Anti-TLE3 | Life Technologies (11372-1-AP) | 1:200 in antibody diluent |
| Anti-TLE5 | Life Technologies (PA5-95534) | 1:500 in antibody diluent |
| Anti-EPHA2 | Cell Signaling (12927S) | 1:200 in antibody diluent |
| Anti-GEMIN4 | Life Technologies (BS-13328R) | 1:500 in antibody diluent |
| Rb Anti-Myc | Cell Signaling (2278S) | 1:1000 in 2% BSA in PBST |

| | | |
|---|---|---|
| Anti-Mouse | LI-COR IRDye 488 nm | 1:1000 in 2% BSA in PBST |
| Anti-Rabbit | LI-COR IRDye 488 nm | 1:1000 in 2% BSA in PBST |

#### Probes used in this study.

| | | |
|---|---|---|
| Biotin-Diazirine | MedChemExpress (HY-154801) | CAS NO. 2845211-64-3 |
| Biotin-Aniline | Millipore Sigma (SML2946) | CAS NO. 769933-15-5 |
| Biotin-Azide | Iris Biotech GmbH (PEG6795.0100) | CAS NO. 2088238-77-9 |
| Biotin-Phenol | Millipore Sigma (SML2135) | CAS NO. 41994-02-9 |
| Biotin-Hydrazide | Chem Scene (CS-6036) | CAS NO. 66640-86-6 |

#### Cloning

For plasmid construction for compartmental studies, backbones from pc3.1 vectors were double-digested and gel purified. Backbones were then fused with gBlock (purchased from IDT) that contains protein target conjugated with a FT tag, or PCR amplified CDS region from commercial plasmid using Phusion polymerase (New England Biolab (NEB), M0536S). Ligated vectors were introduced to NEB 5-alpha competent E. coli cells (NEB, C2987H) by heat shock transformation. For lentivirus construction, lentiviral plasmids pLenti–hPGK-EGFP, pVSV-G, pRSV–Rev, and pMDLg–pRRE were obtained as a kind gift from Dr. J. Burke. The plasmid pLenti-hPGK-H3.1-FT2-FLAG was constructed by PCR amplifying fragment containing H3.1-FT2-FLAG from the pc3.1 vector and fusing the PCR product with digested and linearized pLenti-hPGK backbone. The pMIA plasmids were generated as follows: the murine Runx3 coding sequence (mRUNX3-WT), and RUNX3 fused with TurboID enzyme (mRUNX3-TID) sequences were synthesized and cloned into MSCV-based MIGR1 vector (with GFP or Ametrine reporter) using HiFi DNA Assembly (NEB, #E5520). To conjugate FT tags, oligonucleotides containing FT/FT2 tag were ordered from IDT, self-annealed, and fused with single digested and linearized plasmid at the N-termini of mRUNX3. All ligations were performed using In-Fusion snap assembly kit (Takara Bio, 638947).

3xHA-TurboID-NLS-pCDNA3 was a gift from Alice Ting (Addgene plasmid # 107171 ; http://n2t.net/addgene:107171 ; RRID:Addgene_107171)[1]. Ligated pcDNA3.1(+) vectors were introduced to NEB 5-alpha competent E. coli cells (NEB, C2987H), while lentiviral plasmids were introduced to one shot Stbl3 competent E. coli cells (Thermo Fisher Scientific (Thermo), C737303) by heat shock transformation. Whole plasmid sequencing was performed by Plasmidsaurus using Oxford Nanopore Technology with custom analysis and annotation. For a detailed list of plasmids used in this study refer to supplementary Table 1.

#### HEK293T cell culture and transient transfection

HEK293T cells were cultured in DMEM media (Gibco #11995073) supplied with 10% v/v FBS (R&D Systems S11150), and 100 U ml-1 penicillin 100 µg ml-1 streptomycin (Gibco 15140-122) in 10 cm plates. Cells were maintained in a 37 °C incubator with 5% $CO_2$. For transient transfection, cells grown to 60% confluency in 10 cm plates were treated with 5 µg of plasmid DNA using 15 µL of Lipofectamine 2000 (Thermo, 11668019). Media change was performed 6 hr post transfection, and cells were harvested 24 hr after transfection.

#### Generation of lentiviral particles and stable cell lines

U-2 OS cell line was obtained from Dr. J. Burke as a kind gift. Generation of lentivirus was performed as described previously[2]. Briefly, HEK293T cells with an 80% confluency in a 10 cm dish were co-transfected with 5.85 µg of pLenti-hPGK-H3.1-FT2-FLAG, 1.75 µg of pVSV-G, 1.45 µg of pRSV-Rev, and 2.8 µg of pMDLg-pRRE using 50 µL of Lipofectamine 2000. Medium was replaced 6 hr post-transfection. Medium was collected 24 hr post-transfection and filter-sterilized with a 0.45 µm filter (Thermo, 723-2545). Stable U-2 OS cell line expressing H3.1-FT2-FLAG was generated by incubating U-2 OS cells (24-well plates, 50% confluency) with 0.2 mL of lentivirus particles containing 10 µg/mL of Polybrene (Millipore Sigma (Sigma), TR-1003-G) for 1 hr. Complete DMEM medium was then added to the plate. Selective medium containing 1 µg/mL Puromycin (InvivoGen, ant-pr-1) were replaced 24 hr post-transduction, and cells were maintained in the selective medium for 4 days to allow for complete selection.

#### In vitro screening protocol and western blot analysis

HEK293T cells were lysed with 1X RIPA lysis buffer (RIPA lysis buffer (Sigma, 20-188) diluted in Milli-Q water, supplied with 1x Halt Protease Inhibitor Cocktail EDTA-free (Thermo, 1861279)) and diluted with DPBS buffer (DPBS (10X), no calcium, no magnesium (Thermo, 14200166) diluted in Milli-Q water) to 1 mg/mL concentration validated with Pierce BCA protein assay kit (Thermo, 23227). Cell lysate was mixed well with 2.5 μM of FlAsH-EDT$_2$ and 250 μM probes through vortexing at RT, if no other concentrations have been stated. For quenching tests, 10 mM sodium ascorbate and 5 mM Trolox were added to the mixture prior to irradiation. The mixture was irradiated using with blue or green lamps (LED PAR38 Bulb, 18W (120W equivalent)) for the indicated time. Subsequently, cell lysate was boiled in 1X Laemmli buffer (diluted from 4x Laemmli Sample Buffer (Biorad, 1610747), 2.5 % v/v β-mercaptoethanol) at 95 °C for 10 min and loaded to a pre-cast Bolt™ 4-12% Bis-Tris Plus Mini Protein Gel (Thermo, NW04120BOX). Proteins were transferred to a nitrocellulose membrane (Thermo, 88018) and blocked with 2% Bovine Serum Albumin (BSA) (Fisher Scientific (Fisher), BP9704100) in TBST (diluted from 10x TBST (Bioland Scientific LLC, TBST01-03) with distilled water) for 1 hr at RT. Membranes were then stained with primary antibodies diluted in 2% BSA in TBST at 4 °C overnight on a rocker. Membranes were then washed 3x10 min with TBST and stained with secondary fluorescent antibody diluted in 2% BSA in TBST for 1 hr at RT. Membranes were then washed with TBST for 3x10 min and then imaged on Licor Odyssey CLx scanner. Quantification of the membrane was performed with ImageJ2 (2.14.0).

#### Binding site analysis

Sample preparation and FragPipe analysis were done followed by the protocol stated in[3] with minimal edition. For mass spec sample preparation, purified BSA was dissolved in DPBS in 2 mg/mL concentration, mixed with 2.5 μM of FlAsH-EDT$_2$ (Santa Cruz Biotech Inc, sc-363644) and 250 μM aniline-biotin (An-Bt) (Sigma, SML2946), and irradiated for 2 min under green light. BSA mixed without catalyst but with probe has been used as a negative control. After labeling, samples were desalted with Zeba™ Spin Desalting Columns (Thermo, 89883). The protein sample was reduced with 10 mM 1,4-Dithio-DL-threitol (DTT) (Fisher, 50490514) at 55 °C for 30 min and cysteines alkylated with 22.5 mM Iodoacetamide (IAA) (Fisher, ICN10035125) for 30 min at

RT in dark. Residual IAA was quenched with the addition of 100 mM DTT for 10 minutes at RT. The protein sample was then loaded onto Sera-Mag™ Carboxylate-Modified Magnetic Beads & SpeedBeads (Cytiva, 65152105050250, 45152105050250) for clean-up following established protocols[4]. In brief, 20 µg protein sample were loaded onto 200 µg beads followed by immediate 1:1 v/v dilution of 100% ethanol (Fisher, BP2818500) to initiate protein binding. Following binding, beads were pelleted with magnetic stand, and supernatant was discarded. The pelleted beads were washed three times with 180 µL 80% ethanol in water solution and then resuspended in 40 µL of 100 mM ammonium bicarbonate buffer (Sigma, A6141, diluted with Milli-Q water) containing 1 µg of MS-grade trypsin (Thermo, 25200056) and digested at 37 °C overnight. Digested peptides were bound to new SP3 beads and washed with 100% acetonitrile (MeCN) (Fisher, 501657178) 3x times before eluted with 2% DMSO (Sigma, D8418). Beads were pelleted with a magnetic stand. Samples were eluted and LC-MS/MS analysis of protein digests was carried out using an Orbitrap Fusion Tribrid mass spectrometer (Thermo), following 2 mg capacity ZipTip (Sigma) C18 sample clean-up according to the manufacturer's instructions. Peptides were eluted from an EASY PepMapTM RSLC C18 column (2µm, 100Å, 75µm x 50cm, Thermo), using a gradient of 5-25% solvent B (80/20 acetonitrile/water, 0.1% formic acid) in 45 min, followed by 25-44% solvent B in 15 min, 44-80% solvent B in 0.10 min, a 10 min-hold of 80% solvent B, a return to 5% solvent B in 3 min, and finally with another 3-min hold of 5% solvent B. The gradient was then extended for the purpose of cleaning the column by increasing solvent B to 98% in 3 min, a 98% solvent B hold for 10 min, a return to 5% solvent B in 3 min, a 5% solvent B fold for 3 min, an increase of solvent B to 98% in 3 min, a 98% solvent B hold for 10 min, a return to 5% solvent B in 3 min and a 5% solvent B hold for 3 min and finally, another increase to 98% solvent B in 3 min and a hold of 98% solvent B for 10 min. All flow rates were 250nL/min delivered using a nEasy-LC1000 nano liquid chromatography system (Thermo). Solvent A consisted of water and 0.1% formic acid (FA). Ions were created at 2.5kV using an EASY Spray source (Thermo) held at 50°C. Data dependent scanning was performed by the Xcalibur v 4.0.27.10 software using a survey scan at 120, 000 resolution in the Orbitrap analyzer scanning mass/charge (m/z) 200-2000 followed by higher-energy collisional dissociation (HCD) tandem mass spectrometry (MS/MS) at a normalized collision energy of 30% of the most intense ions at maximum speed, at an automatic gain control of 1.0E4. Precursor ions were selected by the monoisotopic precursor selection (MIPS) setting to peptide and MS/MS was performed on charged species of 2-8 charges at a resolution of

30,000. Dynamic exclusion was set to exclude ions once within a 25 second window. All scan events occurred within a 2-second specified cycle time. The mass spectrometry analysis was performed at The Herbert Wertheim UF Scripps Institute for Biomedical Innovation & Technology, Mass Spectrometry and Proteomics Core Facility (RRID:SCR_023576).

Analysis was performed in FragPipe v21.1. (MSFragger 4.1, IonQuant-1.10.27, philosopher 5.1.0, and DIANN 1.8.2). An open search was done to identify all mass offsets between 300 and 400 Da on the precursor ions, and two mass shifts corresponding to the insertion of aniline-biotin probe have been identified for downstream search. Spectra were searched via LFQ-MBR (IM-MS), with a precursor mass tolerance of 20 ppm and fragment mass tolerance of 20 ppm. Cleavage was specified as strict trypsin with 2 missed cleavages. Modifications selected included cysteine alkylation as a fixed modification, and N-terminal deamidation and methionine oxidation as variable modifications. Spectra were searched against the BSA FASTA obtained from UniProt(Release 202504). Searches were performed for the aniline insertion product mass of either +360.164 or +376.1526 on any residue sites. The results were then filtered for BSA peptides and then analyzed with Perseus (v2.0.7.0), and peptides with an average PSM<2 in the experimental replicates were excluded from the analysis. 2 independent replicates were used per condition. Cutoff was set to be $-\log_{10}$(Adjusted p-value) > 1.3 and $\log_2$(intensity fold change)>1 to identify the enriched peptides with modifications.

#### Cell viability assay

HEK293T cells were transiently transfected to express H3.1-FT2 or mock transfected and were replated in a clear bottom, white walled, 96-well plate (Corning, 3610) and cultured overnight to adhere. The following day, media was removed and replaced with media diluted at indicated concentration and incubated with indicated time. At the end of incubation, media was removed and replaced with complete 1x CellTiter-Fluor reagent (Promega, G6080) diluted in FluoroBrite DMEM (Gibco, A1896701) and incubated for 30 min. Plates were then read 3x for fluorescence on a BioTek Synergy H1 microplate reader. The average of the reads was normalized to DMSO treated wells and fit to a curve using non-linear least squares analysis in Graphpad Prism (Version 10.3.0). For viability comparison between conditions, two-sample *t*-test was performed. Results consist of 3 independent replicating wells per condition.

#### Immunofluorescence assay of labeled cells

Transiently transfected HEK293T cells that express protein conjugated with FT2 tags were plated on pre-treated round coverslips with poly-L-lysine (Sigma, P4707) in 24-well plates. When the cells reached 80% confluency (24 hr post-transfection) they were gently washed with DPBS, then treated with HBSS buffer (Life Technologies, 14025076) containing 2.5 μM FlAsH for 30 min at 37 °C, washed with HBSS containing 125 μM EDT (Sigma, 02390) three times at 37 °C, and habituated for 20 min prior to probe addition. Cells were incubated with HBSS containing 500 μM An-Bt, if no concentration has been stated otherwise, for 30 min to allow for probe penetration, and then irradiated for 5 min if not specifically indicated under green light. Cells were washed with HBSS for three times and incubated for specified time with complete DMEM media ON in a 37 °C incubator with 5% $CO_2$ if longer culturing time was required. Otherwise, cells were then immediately fixed with 4% paraformaldehyde in PBS (Thermo, AAJ61899AP) for 10 min at RT and blocked and permeabilized for 1 hr at RT with 2% BSA in PBST solution (1X DPBS, added with 0.1% Triton Tm X-100 (Fisher, 501781842)). Cells were then stained with primary antibody diluted in 2% BSA in PBST solution at 4 °C ON. Following primary antibody staining, cells were washed with DPBS three times and stained with secondary fluorescent antibody for 1 hr at RT. Coverslips were mounted with mounting media supplied with DAPI (Fisher, NC9524612) and sealed with clear nail polish on glass slides. Slides were imaged on an Olympus FV3000 laser scanning microscope with Abbelight system with a 100X UPLAPOHR OIL NA 1.5 WD 0.12 mm lens, with 405, 488, 561, and 640 nm laser. Colocalization signal analysis was done via ImageJ2.

#### FlAsH-ID labeling for transiently transfected HEK cells

Per replicate, a total of $1.75 \times 10^7$ transiently transfected HEK cells that express protein conjugated with FT tags were harvested 24 hr post-transfection, washed with PBS, and resuspended in 2.5 mL of 2.5 μM FlAsH in HBSS solution with 12.5 μM EDT in 5 mL transparent tube (Fisher, 13-864-406). Cells were incubated in FlAsH solution for 30 min with gentle end-over-end rotation on a tube revolver (Thermo, 9240-11-021) at 37 °C and then pelleted by centrifugation at 400*g* for 4 min at 4 °C. Following 2 washes with HBSS, cells were then consecutively washed with 1 mL of HBSS containing 125 μM EDT 3 times (once for 15 min, and twice for 5 min) at 37 °C with gentle rotation to reduce non-specific FlAsH binding. After washing twice with HBSS buffer to remove excessive EDT, cells were then resuspended in HBSS buffer and rotated for 20 min at RT. Cells were then resuspended in 1 mL of HBSS buffer containing 500 μM An-Bt per $1 \times 10^7$ cells for 30 min at RT to allow probe penetration. Cells were then irradiated with green light in tube for 5 min at 4 °C. The cells were pelleted by centrifugation at 400*g* for 4 min and then washed twice with HBSS buffer to remove excess probe. The washed pellets were then resuspended in 1 mL of RIPA buffer supplied with 0.2% v/v of Pierce™ Universal Nuclease (Thermo, 88702) for 10 min at RT to lyse the cells. Following that, cells were sonicated using a Diagenode Bioruptor Plus for 12 cycles of 30 sec on and 30 sec off at high power at 4 °C. Protein lysate was then clarified through centrifugation at 17,000*g* for 20 min at 4 °C and the protein concentration of the supernatant was determined by BCA assay.

#### TurboID labeling for transiently transfected HEK cells

Optimization and performance of TurboID labeling were accomplished following established protocols[5] with minimal alterations. In brief, per replicate, $3 \times 10^6$ HEK293T cells were plated in a 10-cm plate for transfection the next day. Transient transfection was performed as described above to express RUNX3-TID constructs, while RUNX3-WT with the omittance of the ligase, and TID-NLS, fusion constructs that localize in the nucleus but does not interact with RUNX3, were used as negative controls. 24 hr after transfection, media was replaced with warm biotin-containing (50 μM) complete DMEM media and incubated at 37 °C for 10 min. Labeling was stopped by moving cells onto ice and washed with 5 times of ice-cold PBS. Cells were then harvested by trypsin digestion, pelleted by centrifugation at 400*g* for 4 min, and then lysed with 1 mL of RIPA buffer supplied with 0.2% v/v of Pierce™ Universal Nuclease for 10 min at RT. Following that, cells were sonicated using a Diagenode Bioruptor Plus for 12 cycles of 30 sec on and 30 sec off at high power at 4 °C. Protein lysate was then clarified through centrifugation at 17,000*g* for 20 min at 4 °C and the protein concentration of the supernatant was determined by BCA assay.

#### FlAsH-ID labeling for stable U-2 OS cell line

Per replicate, a total of $1 \times 10^7$ stable U-2 OS cells that express H3.1-FT were harvested, washed with PBS, and resuspended in 2 mL of 2.5 μM FlAsH in HBSS solution with 12.5 μM EDT in 5 mL transparent tube. Cells were incubated in FlAsH solution for 30 min with gentle end-over-end rotation at 37 °C and then pelleted by centrifugation at 400*g* for 4 min at 4 °C. Following 2 washes with HBSS, cells were then consecutively washed with 5 mL of HBSS containing 125 μM EDT 3 times (once for 15 min, and twice for 5 min) at 37 °C with gentle rotation to reduce non-specific FlAsH binding. After washing twice with HBSS buffer to remove excessive EDT, cells were then resuspended in HBSS buffer and rotated for 20 min at RT. Cells were then replated in 10 cm dishes, supplemented with complete DMEM media and incubated ON in a 37 °C incubator with 5% $CO_2$. Cells were transfected with 5 μg of high-molecular-weight poly(I·C) (InvivoGen, tlrl-pic) using 15 μL of Lipofectamine 2000 per 10 cm plate 18 hr post FlAsH treatment. Cells were harvested with trypsinization and resuspended 6 hr after transfection and then resuspended in 5 mL of HBSS buffer containing 500 μM An-Bt per $1 \times 10^7$ cells for 30 min at RT to allow probe penetration. Cells were then irradiated with green light in tube for 5 min at 4 °C. The cells were pelleted by centrifugation at 400*g* for 4 min and then washed twice with HBSS buffer to remove excess probe. The washed pellets were then resuspended in 1 mL of RIPA buffer supplied with 0.2% of nuclease for 10 min at RT to lyse the cells. Following that, cells were sonicated using a Diagenode Bioruptor Plus for 12 cycles of 30 sec on and 30 sec off at high power at 4 °C. Protein lysate was then clarified through centrifugation at 17,000*g* for 20 min at 4 °C and the protein concentration of the supernatant was determined by BCA assay.

For validation of hits with immunofluorescence, U-2 OS cells were plated on pre-treated round coverslips with poly-L-lysine in 12-well plates after FlAsH-ID treatment, and transfected with 0.4 μg of high-molecular-weight poly(I·C) using 1.2 μL of Lipofectamine 2000 per single well 18 hr after treatment. Cells were then incubated with probes and irradiated as described previously and then washed and fixed with 4% PFA in DPBS 6 hr after transfection. Blocking, antibody staining, imaging, and image analysis was performed as described previously.

#### Streptavidin enrichment

Lysate concentration was normalized across all experimental replicates and diluted to 1 mg $mL^{-1}$ with DPBS buffer. Then 1 mL of each sample was incubated with 20 μL of Dynabeads Protein G (Thermo, 10004D) for 30 min at 4 °C with end-over-end rotation to remove non-specific binding proteins. Beads were pelleted by centrifugation at 100*g* for 3 min, and supernatant was incubated with 100 µL of prewashed magnetic Sepharose streptavidin beads (Cytiva Life Sciences, 28985738) overnight at 4 °C with end-over-end rotation. The beads were subsequently washed three times with 1% w/v SDS in DPBS, three times with 1M NaCl in DPBS, and three times with 10% ethanol in DPBS for streptavidin enrichment. For western blot analysis, protein was eluted from beads after washing with PBS for three times by boiling in 1X Laemmli buffer for 15 min under shaking.

#### Label-free proteomics sample preparation

For proteomic samples, the beads were resuspended in 0.5 mL PBS and transferred to a new tube. The beads were washed with 3 × 0.5 mL PBS and 3 × 0.5 mL 100 mM ammonium bicarbonate (Sigma, A6141). The beads were resuspended in 0.5 mL of 3 M urea (Sigma, U5378) in DPBS and 25 µL of 200 mM DTT in 25 mM ammonium bicarbonate was added. The beads were incubated at 55 °C for 30 min. Subsequently, 30 µL of 500 mM iodoacetamide in 25 mM ammonium bicarbonate was added and incubated for 30 min at room temperature in the dark. The supernatant was removed and the beads washed with 3x with 0.5 mL of DPBS and 6x with 0.5 mL of 50 mM ammonium bicarbonate. The beads were resuspended in 0.5 mL of 50 mM ammonium bicarbonate and transferred to a new tube. The beads were resuspended in 40 µL of 50 mM ammonium bicarbonate added with 1.2 µL of trypsin (1 mg/mL in 50 mM acetic acid (Fisher, A11350)), and incubated overnight with end-over-end rotation at 37 °C. After 16 hr, a further 0.8 µL of trypsin was added and the beads were incubated for an extra 1 h at 37 °C. Supernatant was then transferred to new tube and each biological replicate split into two technical replicates. Three biological replicates were used per condition.

#### Label-free proteomics and data analysis

Peptide digests were acidified with TFA to 0.1% (v:v) and desalted using 2 µg capacity ZipTips (Millipore) according to manufacturer instructions. Following drying under vacuum, peptides were re-solubilized in 0.1% formic acid (FA) to a final concentration of 100 ng/µL. Samples were analyzed on a nanoElute 2 (plug-in V2.1.79.0 (1); Bruker) coupled to a Bruker TimsTOF Pro 2 mass spectrometer, equipped with a CaptiveSpray source and a 20 µm zero dead volume (ZDV) Sprayer. Peptides (corresponding to 100 ng) were loaded onto a Thermo Fisher PepMap Neo C18 trap column (300mm X 5mm, 5mm particle size) and then separated on a PepSep Series reverse-phase C18 column (15cm X 150µm, 1.5µm particle size) from Bruker. The column temperature was maintained at 50 °C using an integrated Bruker Column Toaster. The column was equilibrated using 4 column volumes before loading samples in 100% buffer A (99.9% Fisher Optima® LC/MS water, 0.1% FA), with both steps performed at 800 bar. The trap column was equilibrated was equilibrated at 201.6 bar. Samples were separated at 500 nL/min using a linear gradient from 2% to 35% buffer B (99.9% Fisher Optima® LC/MS acetonitrile, 0.1% FA) over 20.0 min before ramping to 95% buffer B (in 0.5 min) and sustained at 95% buffer B for 4.5 min (total separation method time 25.0 min). The Bruker TimsTOF Pro 2 was operated in DIA-PASEF mode using Tims Control v. 5.0.2. Settings for the MS method were as follows: Mass Range 100 to 1700m/z, 1/K0 Start 0.6 V.·/cm2 End 1.4 V·s/cm2, TIMS Ramp and accumulation time 75 ms, Capillary Voltage 1700V, Dry Gas 3 l/min, Dry Temp 200°C, DIA-PASEF settings: 18 MS/MS scans (50m/z windows, 0.21 1/K0 windows, total cycle time 0.74), mass range 300 to 1200, and CID collision energy 20eV (at 0.60, 1/K0) to 65eV (at 1.60, 1/K0). The analysis was performed at The Herbert Wertheim UF Scripps Institute for Biomedical Innovation & Technology, Mass Spectrometry and Proteomics Core Facility (RRID:SCR_023576).

Data was processed via DIANN 1.8.1. Parameters set as follows: trypsin/P digestion, 3 missed cleavages, 3 max. variable modifications, N-term M excision, Ox(M), Ac(N-term) and C carbamidomethylation. Peptide length range was 7-30, precursor charge range 1-4, m/z range 300-1800, and fragment ion range 200-1800. Mass accuracy and MS accuracy were both set to 10. The following settings on the algorithm were checked: “Use isotopologues”, “MBR”, “No shared spectra”, “Heuristic protein inference”. Precursor FDR was set to 1%. A spectral library was used generated via DIANN from all known human proteins (In-Silico spectral library). Resulting matrix.pg file was opened in Perseus. Intensities were inputted as “main”, the rest of the descriptors are categorical. Data was then transformed (log2). Data is annotated by treatment. Normalization was performed via median subtraction. Following this process, a volcano plot was generated utilizing a two-sample *t*-test for statistical significance. If not stated otherwise, Perseus calculated FDR based on *t*-test results was being used to determine the significant hits. For receiver operating characteristic (ROC)-based analysis, lists of true positives (TP) and false positives (FP) were generated from BIOGRID and reported proteome database. Proteins detected in the dataset were ranked by $log_2$(Fold Change) values in descending orders and labeled as TP or FP if matches with the list. At each potential cutoff, TPR and FPR were calculated by dividing the number of TP or FP detected by total number of proteins above the cutoff, and a plot of TPR versus FPR was plotted to show enrichment of TP proteins over FP proteins. The cutoff was determined by using the $log_2$(Fold Change) value that corresponding to the maximum TPR-FPR value. The resulting plots were plotted in R studio (Version 2025.05.1+513) for final figures. Localization of hits were determined using HAS database[6]. STRING was used for gene ontology analyses[7]. BIOGRID database was used to identify interacting protein members[8]. Potential RBPs in the U-2 OS dataset were determined using RBP2GO dataset[9].

#### Isobaric labeling proteomics using tandem mass tags (TMTs)

Following streptavidin-enrichment, the beads were resuspended in 0.5 mL PBS and transferred to a new 1.5 mL Lo-bind tube. Beads were washed with 3× 0.5 mL PBS and 3× 0.5 mL 100 mM ammonium bicarbonate. The beads were resuspended in 0.5 mL of 3 M urea in PBS and 25 µL of 200 mM dithiothreitol in 25 mM ammonium bicarbonate was added. The beads were incubated at 55 °C for 30 min. Subsequently, 30 µL of 500 mM iodoacetamide in 25 mM ammonium bicarbonate was added and incubated for 30 min at room temperature in the dark. The supernatant was removed, and the beads washed 3x with 0.5 mL of DPBS and 6x 0.5 mL of 50 mM Triethylammonium bicarbonate (TEAB) (Thermo, 90114). The beads were resuspended in 0.5 mL of 50 mM TEAB and transferred to a new tube. The beads were resuspended in 40 µL of 50 mM TEAB added with 1.2 µL of trypsin (1 mg/mL in 50 mM acetic acid), and incubated overnight with end-over-end rotation at 37 °C. After 16 hr, a further 0.8 µL of trypsin was added and the beads were incubated for an extra 1 hr at 37 °C. Supernatant was then transferred to new tube and each biological replicate split into two technical replicates.

Meanwhile, TMT-10-plex label reagents (0.8 mg) (Thermo) were equilibrated to room temperature and diluted with 40 µL of anhydrous acetonitrile (Optima grade; 5 min with vortexing) and centrifuged. A total of 20 µL of each TMT reagent was added to the appropriate sample. The reaction was incubated for 2 hr at RT. The samples were quenched with 8 µL of 5% hydroxylamine and incubated for 15 min. The samples were pooled in a new tube and acidified with TFA (16 µL, Optima).

Protein solution samples were desalted using Waters Oasis® OASIS HLB 1cc solid phase extraction cartridges according to the manufacturer's instructions and then dried under vacuum. The dried plex was subsequently solubilized in 400 mL 1M TEAB and fractionated using an Agilent 1100 HPLC system and high pH reversed phase chromatography on a Zorbax Eclipse XDB-C18 column (4.6 x 150mm, 5-micron) from Agilent. Sixty fractions were collected at 0.5 mL/min over 105 min using a gradient of 0-5% solvent B in 10 min, 5-35% solvent B in 60 min, 35-70% solvent B in 15 min, a 10 min hold of 70% solvent B, and finally a return to 5% solvent B in 10 min. Solvent A consisted of 100 mM TEAB and solvent B consisted of ACN. The 60 fractions were dried vacuum, concatenated into 10 fractions, and each peptide fraction was then cleaned up using 2 µg capacity C18 ZipTips (Millipore) according to the manufacturers' instructions. Dried TMT-labelled peptides were reconstituted in 5 µL of 0.1% TFA and on-line eluted into a Fusion Tribrid mass spectrometer (Thermo) from an EASY PepMapTM RSLC C18 column (2µm, 100Å, 75µm x 50cm, Thermo), using a gradient of 6% hold of solvent B (80/20 acetonitrile/water, 0.1% formic acid) for 15 min, followed by 6-28% solvent B in 105 min, then 28-40% solvent B in 10 min, 40-100% solvent B in 10 min, a 10 min hold of 100% solvent B, a return to 5% solvent B in 3 min, and finally a 3 min hold of 5% solvent B. The gradient was then extended for the purpose of cleaning the column by increasing solvent B to 100% in 3 minutes, a 100% solvent B hold for 10 min, a return to 5% solvent B in 3 min, a 5% solvent B fold for 3 min, an increase of solvent B to 100% in 3 min, a 100% solvent B hold for 10 min, a return to 5% solvent B in 3 min and a 5% solvent B hold for 3 min and finally, another increase to 100% solvent B in 3 min and a hold of 100% solvent B for 10 min. All flow rates were 250 nL/min delivered using a Thermo Vanquish Neo UHPLC nano liquid chromatography system. Solvent A consisted of water and 0.1% formic acid. Ions were created at 1.9kV using an EASY Spray source (Thermo) held at 50 °C. A synchronous precursor selection (SPS)-MS3 mass spectrometry method was selected based on the work of Ting et al.[10]. Scans were conducted between 380-2000 m/z at a resolution of 120,000 for MS1 in the Orbitrap mass analyzer at an AGC target of 4E5 and a maximum injection of 50 msec. We then performed CID in the linear ion trap of peptide monoisotopic ions with charge 2-8 above an intensity threshold of 5E3, using a quadrupole isolation of 0.7 m/z and a CID energy of 35%. The ion trap AGC target was set to 1.0E4 with a maximum injection time of 50 msec. Dynamic exclusion duration was set at 60 seconds and ions were excluded after one time within the +/- 10ppm mass tolerance window. The top 10 MS2 ions in the ion trap between 400-1200 m/z were then chosen for HCD at 65% energy. Detection occurred in the Orbitrap at a resolution of 60,000 and an AGC target of 1E5 and an injection time of 120 msec (MS3). All scan events occurred within a 3-second specified cycle time. The analysis was performed at The Herbert Wertheim UF Scripps Institute for Biomedical Innovation & Technology, Mass Spectrometry and Proteomics Core Facility (RRID:SCR_023576).

Raw data was converted to mzmL files using MSConvert(3.0.22216-f3fe83c) following the online tutorial (https://fragpipe.nesvilab.org/docs/tutorial_convert.html). Converted data were then processed using FragPipe using TMT10-MS3 workflow with indicated 229.16293 modification for TMT label adjusted for N-terminus. Samples were filtered by "Reverse" and "Potential contaminant" and were categorized by treatment. FASTA file of UniProt Homo sapiens proteome was included as reference. Protein abundance file was processed through Perseus as previously described, where data were annotated, normalized, and FDR-based *t*-test was performed for statistical significance to generate volcano plots. The resulting volcano plots were plotted and analyzed following the procedures listed in the label-free proteomic workflow.

#### Proximity ligation assay

Transiently transfected HEK293T cells that express protein conjugated with FT tags were plated on pre-treated round coverslips with poly-L-lysine in 24-well plates and fixed with 4% PFA in PBS 24 hr post-transfection. Cells were incubated with pre-chilled 100% MeOH at -20 °C to permeabilize. Once fixed and permeabilized proximity ligation was carried out as recommended by manufacturer using the Duolink in situ red starter kit mouse/rabbit (Sigma-Aldrich DUO92101). In brief, slides were blocked for 1 hr at 37 °C with blocking buffer. Slides were then incubated with primary antibodies diluted with appropriate concentration in antibody diluent buffer at 4 °C ON. Slides were washed with wash buffer A and then incubated with PLA probe plus donkey anti-goat IgG and PLA probe, plus donkey anti-rabbit IgG and PLA probe, or minus donkey anti-mouse IgG antibodies for 1 hr at 37 °C regarding the specific antibody being used. Samples were washed with wash buffer A and incubated for 30 min at 37 °C with Duolink ligase reaction solution. Slides were washed with wash buffer A and incubated for 100 min with duolink amplification solution.

Slides were washed with wash buffer B. Slides were stained with primary Myc-tag antibody ON at 4 °C to identify overexpressed RUNX3, followed by 1 hr of secondary antibody staining at RT. Slides were then mounted with duolink in situ mounting media with DAPI and sealed with clear nail polish for imaging. Slides were imaged on a FV3000 laser microscope as being described previously. Images were acquired from three independent biological replicates, and at least 5 images were taken of each slide in different position. Number of puncta colocalizes with RUNX3 signal were quantified using CellProfiler nuclear speckle counting software. Statistical significance was determined by a one-way ANOVA test followed by post hoc Dunnett's multiple comparisons test of foci per cell using GraphPad Prism with outliers excluded to limit bias from staining inconsistencies.

#### $CD8^+$ T cell transduction

Platinum-E retroviral packaging cells (cultured in DMEM with 4.5 g/L of D-glucose, 10% FBS at 37°C, 5% $CO_2$) were co-transfected with plasmid DNA (see Cloning and supplementary table 1) encoding retroviral constructs and ecotropic packaging helper (pCL-Eco, addgene, #12371) using LT-I TransIT (Mirus, MIR 2304). Retroviral supernatant was collected between 24-48 hr post transfection, and held at 4 °C until use in transduction. $CD8^+$ T cells were isolated from splenocytes using negative selection magnetic bead-based purification according to the manufacturer's protocol (Stemcell EasySep). Purified $CD8^+$ T cells were seeded at a density of $4 \times 10^5$ cells/cm² (final concentration: $0.5 \times 10^6$ cells/mL) onto plates pre-coated with goat anti-hamster capture antibody (50 μg/mL) in T cell Media (DMEM with 4.5 g/L of D-glucose, 10% FBS, supplemented with 1 mM sodium pyruvate, 10 mM HEPES, 1X Gibco MEM NEAA, 1X Gibco MEM Vitamin at 37°C, 5% $CO_2$). Cells were activated using media supplemented with hamster anti-CD3 (clone 2C11) and anti-CD28 (clone 37.51) antibodies (1 μg/mL). Sixteen hours post-activation, culture medium was temporarily removed, and cells underwent spinfection with retroviral supernatant (500 × g, 90 min, 37 °C).

For experiments using transduced $CD8^+$ T cells cultured ex vivo, cells were harvested after 48 hr of anti-CD3/CD28 stimulation and maintained in media containing 10 or 100 U/mL rhIL-2 (NCI). At 6d post cell activation, cells were harvested and used for flow cytometry (Cytek Aurora) and FlAsH-ID labeling. Labeling was performed following the same protocol as described in FlAsH-ID labeling for transiently transfected HEK cells, and data analysis was performed following

Isobaric labeling proteomics using tandem mass tags (TMTs). Antibodies used in this experiment are listed: CD8a (53-6.7), SLAMF6 (13G3), CD103 (2E7), CD62L (MEL-14), PD-1 (29F.1A12), IL-7Rα (A7R34), CD69 (H1.2F3), CD25 (PC61), Lag-3 (C9B7W), Tim-3 (RMT3-23), CD44 (IM7).

#### Mice and $LCMV_{Armstong}$ infection

$CD8^+$ T cells from donor $P14^+dLck\text{-}cre^+Runx3^{+/+}Rosa26\text{-}YFP^{J/J}CD90.1^+CD90.2^-$ and $P14^+dLck\text{-}cre^+Runx3^{fl/fl}Rosa26\text{-}YFP^{J/J}CD90.1^+CD90.2^+$ mice (6 weeks, female) were isolated and transduced with retroviral constructs as described in the previous section. For adoptive transfer, $5\times10^4$ activated cells were injected retro-orbitally into congenically mismatched C57BL/6 mice (6 weeks, female). 1 hr after adoptive transfer, recipients were infected with $2\times10^5$ PFU of $LCMV_{Arm}$ resulting in an acute infection.

Peripheral blood was collected at 8, 15, and 30 days post-infection (p.i.) into heparinized tubes, subjected to red blood cell lysis, washed, and stained with fluorochrome-conjugated antibodies against CD8α (53-6.7), Thy1.1 (OX-7), Thy1.2 (30-H12), CX3CR1 (SA011F11), KLRG1 (2F1/KLRG1), CD62L (MEL-14), CD27 (LG.3A10), IL-7Rα (A7R34), SLAMF6 (13G3), and CXCR3 (CXCR3-173). Samples were acquired on a Cytek Aurora flow cytometer. Spleens were harvested at 30 p.i., processed into single-cell suspensions, subjected to red blood cell lysis, and stained using the same antibody panel.

#### Runx3 and Arid1a bioinformatics analysis

*Runx3 ChIP:* Single-ended FASTQ files (GSE50131) were trimmed and aligned to GRCm39 with Bowtie2. Peaks were called for the naïve and 6d *ex vivo* samples with MACS2 callpeak (p-value < 0.01), and blacklist regions filtered. Runx3-bound peaks were defined as those present in the 6d *ex vivo* sample (relative to input DNA) but absent in the naïve consensus set using subsetByOverlaps(…,invert=TRUE) from the GenomicRanges package in R.

*Arid1a CUT&RUN:* Paired-end FASTQ files (GSE228380) were analyzed similarly to Runx3 ChIP-seq data, with the exception of duplicate removal using samtools markdup, and setting the IgG control sample as control during MACS2 peak calling.

*ATAC-seq:* Paired-ended reads (GSE111149 and GSE228171) were trimmed, aligned to GRCm39 and duplicates removed using samtools markdup. Reads were shifted to account for Tn5 insertion bias and peaks were called with MACS2 callpeak (--nomodel --shift -100 --extsize 200).

HOMER findMotifsGenome.pl was used to quantify enrichment of transcription factor motifs within filtered peak subsets, and enrichment ratio (target / background) and p-values for each motif were used to make enrichment plots.

To assign genes to Runx3/Arid1a co-bound peaks (whose accessibility was instigated by T cell activation), a window-based candidate gene assignment algorithm was implemented in Python. For a symmetric window size W (set to 100,000 base pairs), each peak $p_i$ defines an extended interval:

$$p_i^{(W)} = c_i, s_i - W, e_i + W$$

Where $c_i$, $s_i$, and $e_i$ represent the chromosome number, start and end genomic co-ordinates of $p_i$. A gene $g_j$ is considered "proximal" to $p_i$ if $p_i^{(W)}$ and $g_j$ overlap by at least 1 bp. Each genes final score is computed by aggregating contributions from all proximal peaks:

$$S_j = \sum_{i=1}^{N} prox(p_i, g_j)$$

#### References

1. Branon, T. C. *et al.* Efficient proximity labeling in living cells and organisms with TurboID. *Nat. Biotechnol.* **36**, 880–887 (2018).

2. Burke, J. M., Moon, S. L., Matheny, T. & Parker, R. RNase L Reprograms Translation by Widespread mRNA Turnover Escaped by Antiviral mRNAs. *Mol. Cell* **75**, 1203-1217.e5 (2019).

3. Huth, S. W. *et al.* μMap Photoproximity Labeling Enables Small Molecule Binding Site Mapping. *J. Am. Chem. Soc.* **145**, 16289–16296 (2023).

4. Hughes, C. S. *et al.* Ultrasensitive proteome analysis using paramagnetic bead technology. *Mol. Syst. Biol.* **10**, 757 (2014).

5. Cho, K. F. *et al.* Proximity labeling in mammalian cells with TurboID and split-TurboID. *Nat. Protoc.* **15**, 3971–3999 (2020).

6. Thul, P. J. & Lindskog, C. The human protein atlas: A spatial map of the human proteome. *Protein Sci.* **27**, 233–244 (2018).

7. Szklarczyk, D. *et al.* STRING v11: protein-protein association networks with increased coverage, supporting functional discovery in genome-wide experimental datasets. *Nucleic Acids Res.* **47**, D607–D613 (2019).

8. Oughtred, R. *et al.* The BioGRID database: A comprehensive biomedical resource of curated protein, genetic, and chemical interactions. *Protein Sci.* **30**, 187–200 (2021).

9. Caudron-Herger, M., Jansen, R. E., Wassmer, E. & Diederichs, S. RBP2GO: a comprehensive pan-species database on RNA-binding proteins, their interactions and functions. *Nucleic Acids Res.* **49**, D425–D436 (2021).

10. Ting, L., Rad, R., Gygi, S. P. & Haas, W. MS3 eliminates ratio distortion in isobaric multiplexed quantitative proteomics. *Nat. Methods* **8**, 937–940 (2011).

### Supplementary Tables

#### Supplementary Table 1-4

Supp. Table 1-4 are provided as XL files and contain all plasmid sequences used in the study and proteomics data.

### Uncropped western blots

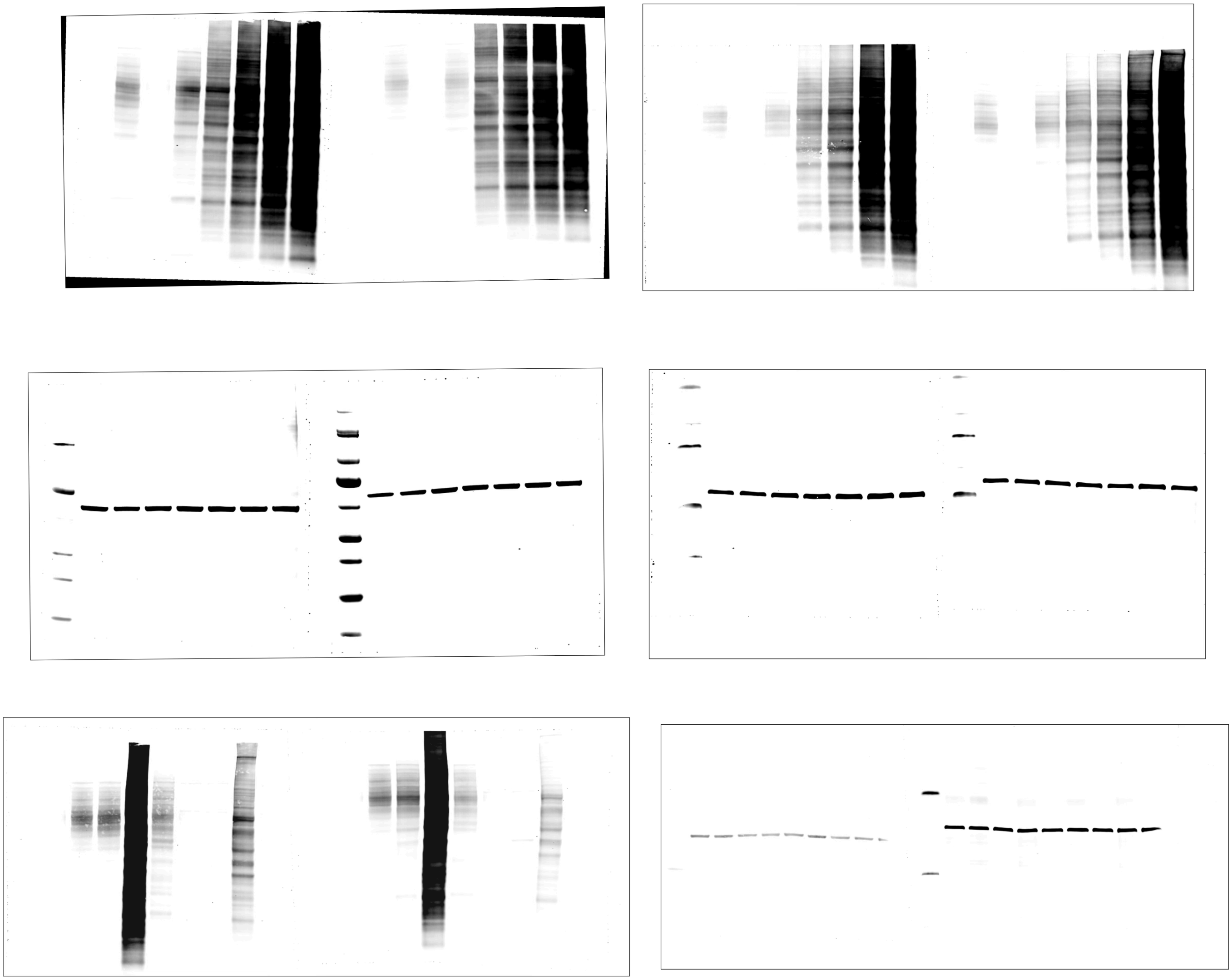

Uncropped blot from Supplementary Fig.1

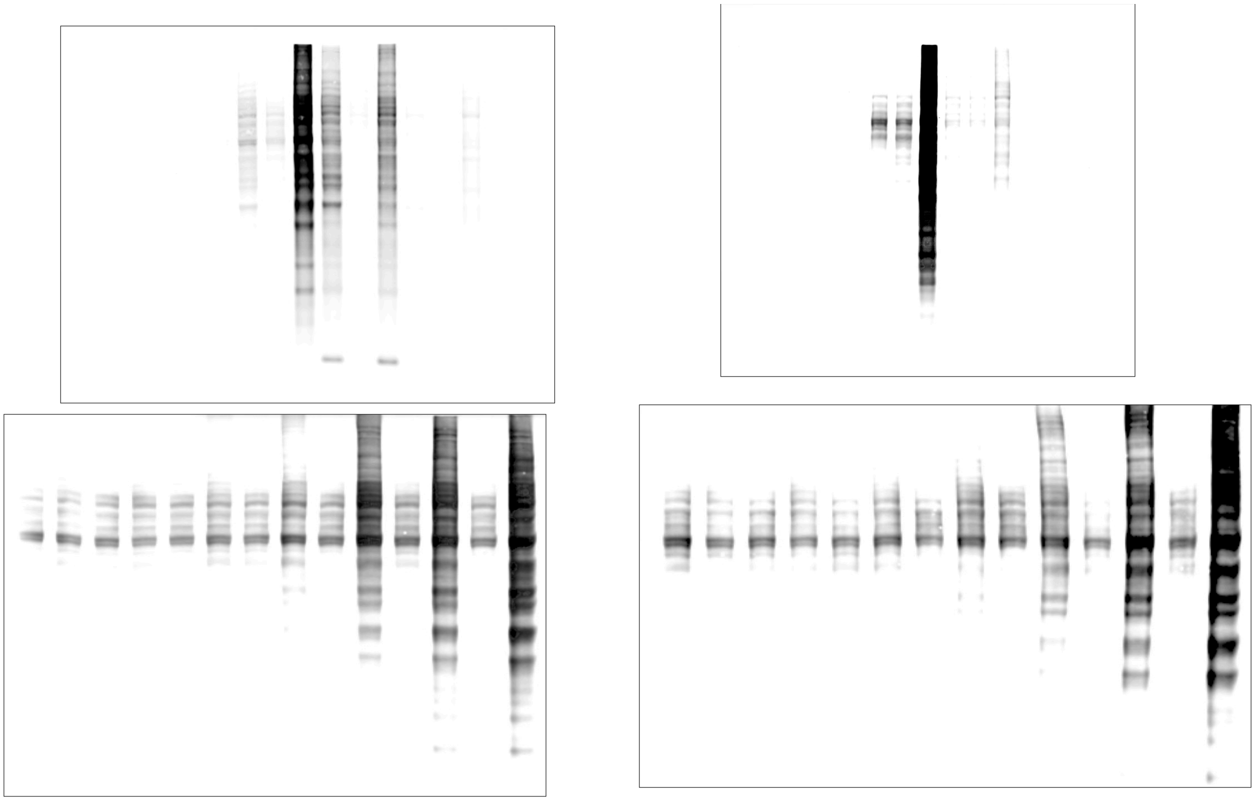

Uncropped blot from Fig. 1

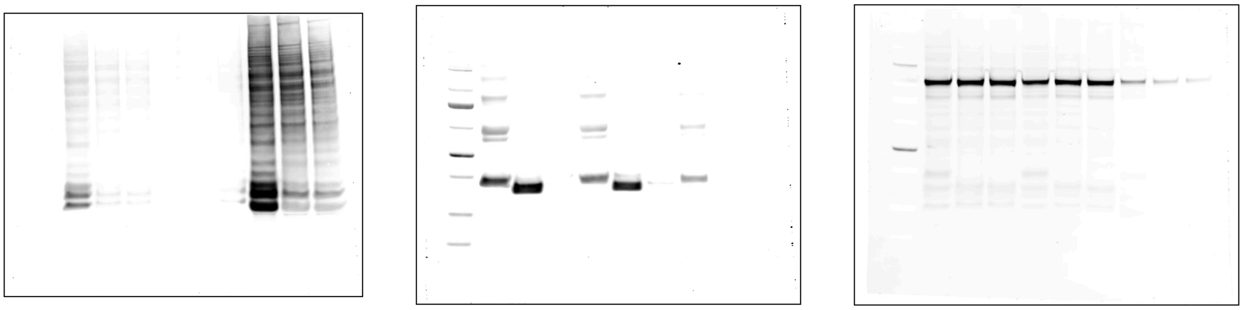

Uncropped blot from Fig. 2
